# The exon junction complex regulates the release and phosphorylation of paused RNA polymerase II

**DOI:** 10.1101/271544

**Authors:** Junaid Akhtar, Nastasja Kreim, Federico Marini, Giriram Kumar Mohana, Daniel Brune, Harald Binder, Jean-Yves Roignant

## Abstract

Promoter proximal pausing of RNA polymerase II (Pol II) is a widespread transcriptional regulatory step across metazoans. Here we find that the nuclear exon junction complex (pre-EJC) plays a critical and conserved role in this process. Depletion of pre-EJC subunits leads to a global decrease in Pol II pausing and to premature entry into elongation. This effect occurs, at least in part, via non-canonical recruitment of pre-EJC components at promoters. Failure to recruit the pre-EJC at promoters results in increased binding of the positive transcription elongation complex (P-TEFb) and in enhanced Pol II release. Notably, restoring pausing is sufficient to rescue exon skipping and the photoreceptor differentiation defect associated with depletion of pre-EJC components *in vivo*. We propose that the pre-EJC serves as an early transcriptional checkpoint to prevent premature entry into elongation, ensuring proper recruitment of RNA processing components that are necessary for exon definition.

## HIGHLIGHTS

- Depletion of pre-EJC components leads to widespread transcriptional changes in fly and human cells
- Pre-EJC components associate at gene promoters via binding to Pol II and nascent RNAs
- The pre-EJC stabilizes Pol II pausing at least in part by restricting P-TEFb binding
- The pre-EJC regulates exon definition by preventing premature release of Pol II.

## INTRODUCTION

Transcripts produced by RNA polymerase II (Pol II) undergo several modifications before being translated, including 5’-end capping, intron removal, 3’-end cleavage and polyadenylation. These events usually initiate co-transcriptionally while the nascent transcript is still tethered to the DNA by Pol II (Brugiolo et al., 2013; Girard et al., 2012; Khodor et al., 2011; Tilgner et al., 2012). This temporal overlap is important for the coupling between these processes (Bentley, 2014; Custodio and Carmo-Fonseca, 2016; Herzel et al., 2017; Jonkers and Lis, 2015; Saldi et al., 2016). Initially, Pol II is found in a hypophosphorylated form at promoters. At the onset of initiation, the CTD of Pol II becomes phosphorylated at the Ser5 position. Pol II subsequently elongates and often stalls 20-60 nucleotides downstream of transcription start sites (TSS), an event commonly referred to promoter proximal pausing (Gilmour and Lis, 1986; Rougvie and Lis, 1988). Promoter proximal pausing of Pol II is widely seen at developmentally regulated genes, and is thought to play critical roles in facilitating rapid and synchronous transcriptional activity upon stimulation (Adelman and Lis, 2012; Gaertner et al., 2012; Kwak and Lis, 2013; Levine, 2011; Liu et al., 2015; Smith and Shilatifard, 2013). Pol II pausing is also suggested to act as a checkpoint influencing downstream RNA processing events such as capping and splicing, but evidence for this function is still limited. The transition from the paused state to elongation is promoted by the positive transcription elongation factor (P-TEFb) complex, which includes the cyclin-dependent kinase 9 (Cdk9) and cyclin T (Gressel et al., 2017; Luo et al., 2012; Marshall and Price, 1995; Peterlin and Price, 2006). P-TEFb phosphorylates Ser2 of the CTD as well as the negative elongation factor (NELF) and DRB sensitivity-inducing factor (DSIF), leading to the release of Pol II from promoter (Fujinaga et al., 2004; Marshall et al., 1996; Ni et al., 2008). Another related kinase, Cdk12, was also recently suggested to affect Pol II pausing after its recruitment through Pol II-associated factor 1 (PAF1) (Chen et al., 2015; Yu et al., 2015)

The exon junction complex (EJC) is a ribonucleoprotein complex, which assembles on RNA upstream of exon-exon boundaries as a consequence of pre-mRNA splicing (Le Hir et al., 2000a; Le Hir et al., 2000b). The spliceosome-associated factor CWC22 is essential to initiate this recruitment (Alexandrov et al., 2012; Barbosa et al., 2012; Steckelberg et al., 2015; Steckelberg et al., 2012). The nuclear EJC core complex, also called pre-EJC, is composed of the DEAD box RNA helicase eIF4AIII (Shibuya et al., 2004), the heterodimer Mago nashi (Mago) (Kataoka et al., 2001) and Tsunagi (Tsu/Y14) (Hachet and Ephrussi, 2001; Kim et al., 2001). The last core component, Barentsz, joins and stabilizes the complex during or after export of the RNA to the cytoplasm (Degot et al., 2004). The EJC has been shown to play crucial roles in post-transcriptional events such as RNA localization, translation and nonsense-mediated decay (Boehm and Gehring, 2016; Gerbracht and Gehring, 2018; Le Hir et al., 2016). These functions are mediated by transient interactions of the core complex with effector proteins (Tange et al., 2005).

Previous studies have identified an additional role for the pre-EJC in pre-mRNA splicing (Hayashi et al., 2014; Malone et al., 2014). In absence of the pre-EJC, many introns containing weak splice sites are retained. The pre-EJC facilitates removal of weak introns by a mechanism involving its prior deposition to adjacent exon junctions. This intron definition activity is mediated via the EJC splicing subunits RnpS1 and Acinus through a mechanism that is not yet fully understood (Hayashi et al., 2014; Malone et al., 2014). Other studies demonstrated an additional role for the pre-EJC in exon definition in both *Drosophila* and human cells (Ashton-Beaucage et al., 2010; Roignant and Treisman, 2010; Wang et al., 2014). In *Drosophila*, loss of Mago in the eye leads to several exon skipping in *MAPK*, resulting in photoreceptor differentiation defects. Other large transcripts, often expressed from heterochromatic regions, show the same Mago-splicing dependency. Similarly, in human, exons flanked by longer introns are more dependent on the EJC for their splicing. Interestingly, this exon definition activity seems mostly independent of EJC splicing subunits (Wang et al., 2014).

Here, we investigated the mechanism underlying the role of the pre-EJC in exon definition in *Drosophila*. We observed that depletion of pre-EJC components, but not of the EJC splicing subunit RnpS1, led to genome-wide changes in the phosphorylation state of Pol II and to a global decrease in promoter proximal pausing. This change in phosphorylation state is concomitant with changes in histone modifications and chromatin accessibility. In addition, we found that pre-EJC components associate with promoter regions, providing a link between the pre-EJC and the transcription machinery. Co-immunoprecipitation experiments indicate that Mago associates with Pol II but this association is largely dependent on nascent RNA. Upon knockdown (KD) of pre-EJC components, Cdk9 binding to Pol II is increased, partly accounting for the premature Pol II release. Remarkably, genetically reducing Pol II pausing rescues exon skipping events and the eye phenotype associated with the KD of pre-EJC components, indicating that restraining Pol II release into gene bodies is sufficient to complement the loss of pre-EJC components *in vivo*. Altogether, our results demonstrate a direct role for the pre-EJC in controlling the transcriptional state. Our data support a mechanism in which the pre-EJC facilitates the definition of exons surrounded by large introns by maintaining a timely transition into transcription elongation via the control of promoter proximal pausing of RNA Pol II.

## RESULTS

### 1 Pre-EJC components Regulate Expression and Splicing of Large Intron-containing Genes

Previous studies demonstrated a critical role for Mago in preventing exon skipping in *MAPK* and other large intron-containing transcripts in *Drosophila* cells (Ashton-Beaucage et al., 2010; Roignant and Treisman, 2010). To address whether this function was shared with other EJC components we depleted each subunit of the core complex as well as the associated splicing subunit RnpS1 (Figure S1A). We found that depletion of pre-EJC components (eIF4AIII, Mago and Y14) more dramatically affects gene expression compared to the depletion of the cytoplasmic subunit Btz and RnpS1 (Figure S1B). As previously shown for Mago depletion, large intron-containing transcripts were preferentially affected upon KD of pre-EJC subunits, while depletion of Btz or RnpS1 did not show this trend (Figure S1C). This reduced expression level correlates with an overrepresentation of exon skipping observed in these large intron-containing genes, such as *MAPK* (Figures S1D-E). This was specific to Mago depletion, as knockdown of Mago in S2R+ cells transfected with RNAi resistant Mago cDNA did not result in splicing defect of *MAPK* (Figure S1F). Furthermore, the loss of pre-EJC resulted in higher degree of exon skipping events, than its cytoplasmic component Btz or accessory subunit RnpS1 (Figure S1G). Altogether these results indicate that the pre-EJC is required for proper expression and splicing of large intron-containing genes.

### 2 Depletion of pre-EJC factors Leads to a Genome-Wide Alteration of Pol II Phosphorylation

Introns are being spliced while nascent RNA is still tethered to Pol II, allowing coupling between splicing and transcription machineries (Braunschweig et al., 2013; Custodio and Carmo-Fonseca, 2016; Herzel et al., 2017; Moehle et al., 2014; Naftelberg et al., 2015; Saldi et al., 2016). To address the possibility that the pre-EJC regulates splicing via modulation of transcription, we performed chromatin immunoprecipitation (ChIP) experiments using chromatin extracts from *Drosophila* S2R+ and antibodies that specifically recognize the different forms of Pol II. We first interrogated by qPCR the *MAPK* locus, as splicing of its pre-mRNA is strongly affected upon depletion of pre-EJC components (Figures S1D and S1E, (Ashton-Beaucage and Therrien, 2011; Roignant and Treisman, 2010). Interestingly, we found that Mago KD results in decrease of Pol II occupancy at the 5’ end of *MAPK* while the distribution in the rest of the gene body was comparable to the control (Figure 1A). This was specific to Mago depletion as reintroducing Mago cDNA restores the wild type profile (Figure S2A). The Ser2 phosphorylated form of Pol II also exhibited a distinct pattern. While its level mildly decreases at the transcription start site (TSS), it significantly increases along the gene body (Figure 1A). To examine whether these changes in Pol II distribution were widespread, we performed ChIP-Seq experiments in control and Mago KD conditions. To exclude the possibility of changes in Pol II occupancy driven by differences in immunoprecipitation efficiency and technical variance during library preparation in different knock down conditions, we used yeast chromatin as “spike-in” control (Orlando et al., 2014). With this approach, we confirmed the decrease in Ser2P levels and Pol II at the promoter region and an increase within the gene body of *MAPK* (Figure 1B). Examining Pol II and Ser2P profiles in a genome-wide manner show extensive changes with decrease at the TSS and increase towards transcription end sites (TES) (Figures 1C-E). Similar changes in Pol II occupancy were observed upon depletion of Y14 and eIF4AIII (Figures S2B-E) but neither of RnPS1 nor of Btz (Figures S2E-G). Thus, these results demonstrate that pre-EJC components regulate Pol II distribution.

**Figure 1.**
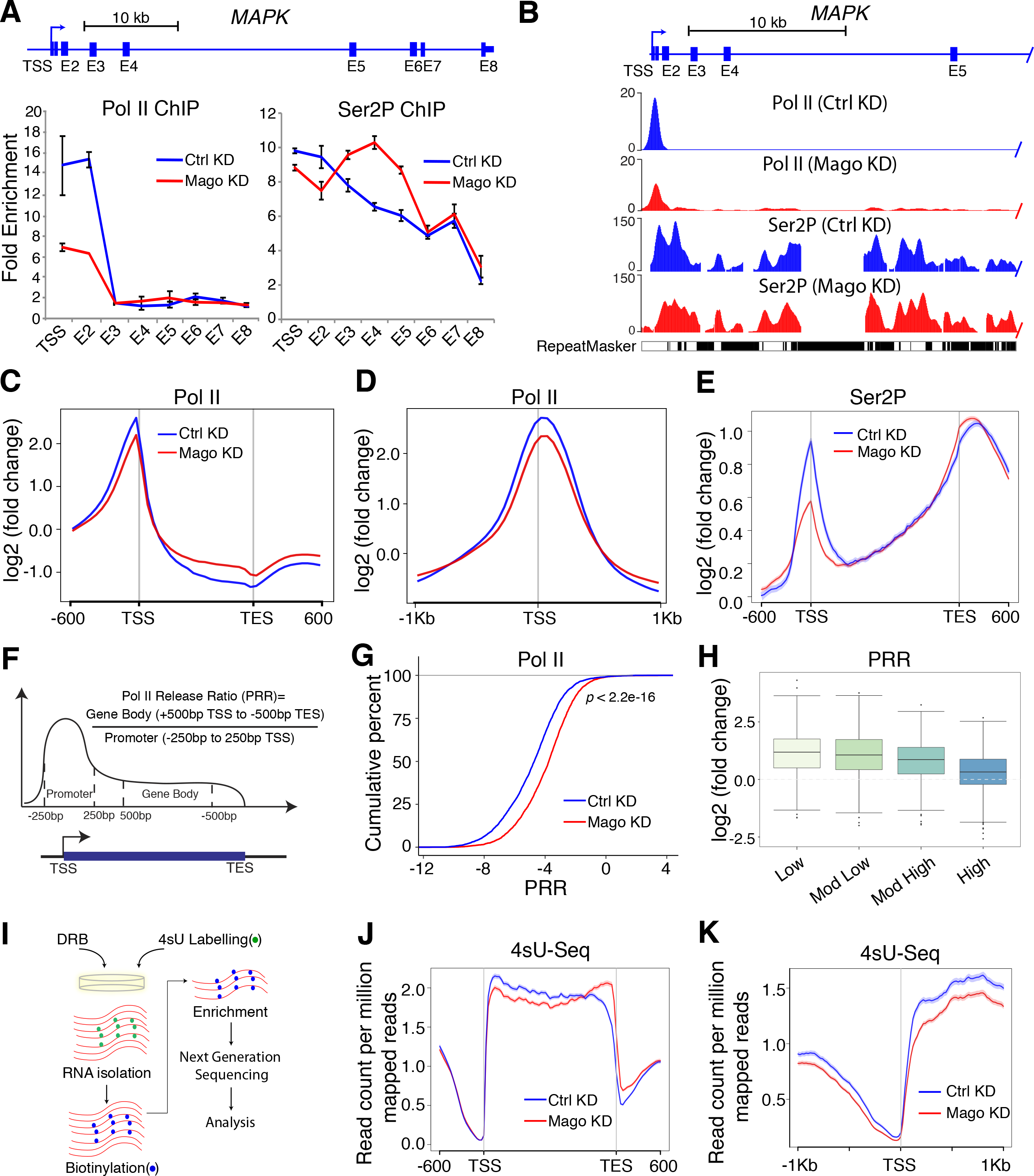
Mago Prevents Premature Release into Transcription Elongation. **(A)** ChIP-qPCR analysis of Pol II and Ser2P occupancies at *MAPK* locus. The tested regions for enrichment are shown in the scheme. Error bars indicate the standard deviation from the mean in three biological replicates. **(B)** Track examples of total Pol II and Ser2P ChIP-Seq from S2R+ cells extracts, after either control or Mago knockdown. The tracks are average of two independent biological replicates after input and ““spike-in”” normalization. Shown here are the profiles on *MAPK*, a well-described pre-EJC target gene. **(C, D)** Averaged metagene profiles of total Pol II occupancies from two independent biological replicates after ““spike-in”” normalization in control and Mago-depleted cells, −600bp upstream of transcription start sites (TSS) and +600bp downstream of transcription end sites (TES) **(C)** or centered at the TSS in a ± 1 Kb window **(D)**. Log2 fold changes against input control are shown on Y-axis, while X-axis depicts genomic coordinates. **(E)** Averaged metagene profiles of two independent biological replicate of Ser2P occupancies in control and Mago-depleted cells. Log2 fold changes against input control are shown on Y-axis, while X-axis depicts scaled genomic coordinates. **(F)** Schematic representation of the calculation of the Pol II release ratio (PRR). The promoter is defined as 250bp upstream and downstream of TSS, while the gene body is 500bp downstream of TSS to 500bp upstream of transcription end site (TES). **(G)** The empirical cumulative distribution function (ECDF) plot of computed PRR in control and Mago knockdown conditions, after “spike-in” normalization. *P*-value is derived from two-sample Kolmogorov-Smirnov test. **(H)** Box plots showing changes in PRRs upon Mago depletion when compared to control, separated into different PRR quartiles. **(I)** Schematic depiction of the DRB-4sU-Seq approach. **(J)** Metagene profile of nascent RNA from non-DRB treated 4sU-Seq data in control and Mago-depleted cells. Averaged read counts per million of mapped reads of two independent biological replicates from 4sU-Seq are shown on Y-axis while X-axis depicts scaled genomic coordinates. **(K)** Metagene profile of nascent RNA from non-DRB treated 4sU-Seq data in control and Mago-depleted cells, centered at the TSS in a ± 1 Kb window. Nascent RNA was fragmented to ≤ 100bp during enrichment. Averaged read counts per million of mapped reads of two independent biological replicates from 4sU-Seq are shown on Y-axis while X-axis depicts genomic coordinates.

### 3 Pre-EJC Components Facilitate Promoter Proximal Pausing of RNA Pol II

To further investigate the changes of the transcriptional state in pre-EJC-depleted cells, we compared the Pol II release ratio (PRR) in control versus KD conditions for pre-EJC components. To this end, we calculated the ratio of Pol II occupancy between gene bodies and promoter regions (Figure 1F). Notably, we found that the PRR was significantly higher upon depletion of pre-EJC components compared to control conditions (Figures 1G and S2H). Changes in the PRR were specific to pre-EJC depletion, as depletion of RnpS1 did not result in similar alteration (Figure S2I). These results indicate a specific role for pre-EJC components in controlling transcription by regulating promoter proximal pausing of RNA Pol II. We next divided the changes in PRR upon Mago KD into four equal size quartiles, from low to high. When classified accordingly, the quartile with the lowest PRRs showed highest change in PRR upon Mago KD (Figure 1H). This indicates that strongly paused genes are more affected upon the loss of Mago.

To conclusively determine the changes in PRR upon pre-EJC depletion, we carried out a 4sU-seq approach similar to the method recently employed for sequencing all transient transcripts in human cells (transient transcriptome sequencing (TT-Seq), (Schwalb et al., 2016)), with some modifications (see material and methods) (Figure 1I). The resulting 4sU-Seq metagene profile revealed lower read counts at the TSS upon Mago depletion, while an increase towards the 3’ end of transcripts was observed, confirming our previous Pol II ChIP-Seq findings (Figures 1J and 1K). We next coupled this approach with a treatment of the reversible compound 5,6-dichlorobenzimidazole 1-β-D-ribofuranoide (DRB), to monitor nascent transcription overtime (4sUDRB-seq) (Singh and Padgett, 2009). We calculated the elongation speed of all genes longer than 10 kb, as this approach is only amenable for large genes. We divided the genes that passed this size criterion into multiple similar sized bins and examined the progression of signal of the nascent RNA over time as described earlier (Figures S3A and S3B) (Fuchs et al., 2014). This analysis revealed that the average elongation rate in *Drosophila* S2 cells is slower in comparison to human. We found an average rate of 1 kb per minute, contrasting with the 3 to 4 kb per minute reported in human cells (Figure S3C). Nonetheless, our calculation is in agreement with the rate that was previously measured on a few individual *Drosophila* genes (Ardehali and Lis, 2009; O’Brien and Lis, 1993; Thummel et al., 1990; Yao et al., 2007). Furthermore, in contrast to the widespread change in promoter proximal pausing, we did not observe a significant alteration of the average elongation rate in Mago-depleted cells (Figure S3C). The moderate gene-to-gene variation of the elongation rate does not correlate with changes in exon inclusion (Figure S3D), suggesting that Mago does not control exon definition via the regulation of Pol II elongation kinetics. Altogether, our data indicate that premature Pol II release has little effect on the speed of Pol II elongation and, conversely, that the global reduction of Pol II pausing observed upon Mago KD is not a consequence of an increase in the elongation rate.

### 4 Pre-EJC components Associate at Promoter Regions to Regulate Pol II Pausing

We next aimed to address the mechanism by which the pre-EJC controls Pol II pausing. We wondered whether the global effect on Pol II pausing results from misexpression of a critical factor involved in this process. To answer this question, we re-analyzed our transcriptome dataset to search for potential targets involved in promoter proximal pausing or in Pol II release. However, none of the known pausing factors, including Cdk9, Spt5, subunits of the NELF complex, GAGA, Med26, TFIID, exhibit altered expression upon KD of pre-EJC components (Figures S4A-D). The absence of obvious candidates prompted us to test whether the pre-EJC may directly associate with chromatin to regulate Pol II pausing. We then performed ChIP-Seq experiments using cells expressing HA-tagged versions of EJC components. Strikingly, we found that pre-EJC components associate with chromatin, which was in contrast to RnpS1 (Figures 2A and S5A). We found that most of the enriched binding regions fall at promoters of expressed genes (Figure 2B and S5B). Example tracks are shown in Figures 2C and S5C. These results are consistent with a recent study showing association of Y14 to promoters in *Drosophila* cells (Choudhury et al., 2016). The degree of overlap between the bound targets of pre-EJC components was relatively high (34%), indicating that they have numerous target genes in common (n=756) (Figure S5D). Importantly, enrichment of Mago-HA was strongly decreased if cells were subjected to Mago KD, demonstrating the specificity of the signal (Figure S5B).

**Figure 2.**
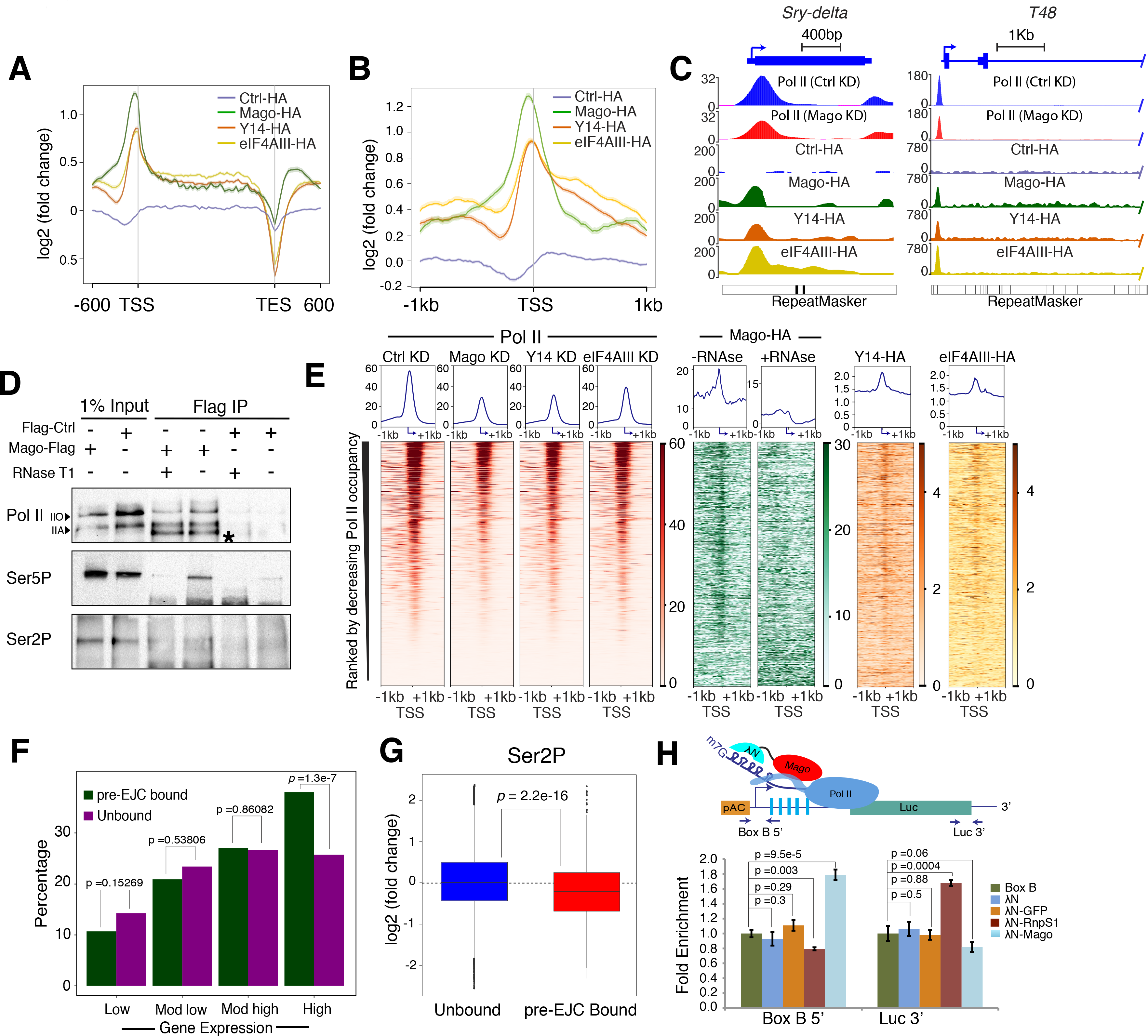
Mago Binding to Promoter Regions Modulates Pol II Pausing. **(A)** Averaged metagene profiles from two independent biological replicates of ChIP-Seq performed with HA-tagged Mago, Y14, eIF4AIII and Ctrl. Log2 fold changes against input control are shown on Y-axis while X-axis depicts scaled genomic coordinates. **(B)** Averaged metagene profiles from two independent biological replicates of ChIP-Seq performed with HA-tagged Mago, Y14, eIF4AIII and Ctrl, centered at the TSS in a ± 1 Kb window. Log2 fold changes against input control are shown on Y-axis while X-axis depicts genomic coordinates. **(C)** Input normalized and replicate averaged track examples of ChIP-Seq experiments from S2R+ cell extracts transfected with HA-tagged Mago, Y14, eIF4AIII, or Ctrl. Shown here is recruitment of pre-EJC components to an intronless (*Sry-delta*) and intron-containing (*T48*) genes **(D)** Co-immunoprecipitation using anti-Flag antibody from cell extracts expressing either Flag-Mago or Flag alone (Flac-Ctrl), revealed with total Pol II, Ser5P and Ser2P antibodies. Note that Mago interacts with total Pol II, both hypo (IIA) and hyper phosphorylated (IIO) forms (indicated with the arrowheads). Mago also interacts with Ser5P but not with Ser2P, and this interaction with Pol II is partially dependent on RNA. **(E)** Heatmaps of HA-tagged pre-EJC components and total Pol II, centered at the TSS (−1kb to +1kb). Rows indicate all the genes bound by Pol II and are sorted by decreasing Pol II occupancy. The color labels to the right indicate the levels of enrichment. **(F)** Histogram showing percentage of pre-EJC bound genes amongst different quartiles of genes expressed in control condition. For quartile classification, all of the expressed genes in S2R+ cells were divided into four equal sized quartiles according to the level of expression, from low to high level. *P*-values for significance of the associations, derived from Fisher’s exact test, are shown on top of the histogram. **(G)** Log2 fold changes in Ser2P level at the TSS upon control and Mago knockdowns, when separated according to pre-EJC binding. *P*-value is derived from a two-sample t-test. **(H)** Recruitment of Mago at the 5’ end of RNA is sufficient to induce pausing. (Top) Schematic of the BoxB-XN tethering assay. BoxB sequences (blue rectangles) were inserted upstream of the CDS of Firefly luciferase (green rectangle). The λN peptide (blue) was fused to Mago (shown in red), GFP or RnpS1, and transfected into S2R+ cells along with the modified Firefly luciferase plasmid as well as with a Renilla luciferase construct. (Bottom) Quantification of the ChIP experiment. Chromatin was prepared for the different conditions and followed by immunoprecipitation using antibody directed against total Pol II. The enrichment of Pol II at the promoter and at the 3’ end of Firefly luciferase was calculated after normalizing against a negative loci and Renilla. The enrichment for three independent biological replicates is shown along with *P*-values, for tested conditions.

To get further insights into the mechanisms underlying the association of pre-EJC components at promoters, we tested whether Mago could associate with RNA Pol II. Co-immunoprecipitation experiments using Flag-tagged Mago revealed mild but reproducible interaction with Pol II (Figure 2D). Importantly, we found that Mago interacts with Pol II Ser5P but fails to associate with Ser2P, potentially explaining the enrichment of pre-EJC binding at promoter regions. In addition, interaction with Pol II was reduced after treatment with RNase T1, indicating that a RNA intermediate facilitates this association (Figure 2D). In agreement with this observation, most of Mago binding to promoters was lost when the chromatin was treated with RNAse T1 prior to immunoprecipitation (Figures 2E and S5A, S5B). Furthermore, RNA immunoprecipitation experiments confirm binding to RNA, even to intronless transcripts, indicating that in contrast to canonical EJC deposition, association of Mago at promoters occurs independently of pre-mRNA splicing (Figure S4E). This was further confirmed by the absence of requirement for the spliceosome-associated factor CWC22 for Mago binding (Figures S4F and S4G) as well as the modest effect of a splicing inhibitor, in particular at intronless-bound genes (Figure 4G). Finally, our ChIP-qPCR experiments revealed that cells treated with α-amanitin for seven hours before cross-linking show reduced Mago binding, implying the involvement of nascent RNA for pre-EJC binding (Figure S4H). Collectively our data indicate that pre-EJC components associate at promoters via binding to Pol II, which is phosphorylated on Ser5, and strongly suggest that nascent RNA is required to stabilize this interaction.

### 5 Pre-EJC binding to Nascent RNA is Sufficient to Increase Pausing

In order to establish potential relationship between pre-EJC-bound genes and promoter proximal pausing, we compared pre-EJC binding (genes bound by all pre-EJC components, n=756) with several criteria. First, heatmaps show a positive correlation between Pol II and pre-EJC binding (Figure 2E). Consistently, the proportion of Mago or pre-EJC-bound genes was higher on highly expressed genes (Figures S2F and S5E). Second, we noticed that binding was also enriched at genes that have a low PRR in wild type condition, or in other words, that are strongly paused (Figure S5F). Lastly, we found a positive correlation between pre-EJC binding and changes in Ser2P levels upon Mago KD (Figure 2G, *P* < 2.2 × 10^−16^). Altogether these results indicate possible interplay between pre-EJC binding and modulation of promoter proximal pausing.

To firmly address a direct role of the pre-EJC in the regulation of Pol II pausing we tethered Mago to the 5’ end of nascent RNA and performed ChIP-qPCR experiments to evaluate the effect on Pol II occupancy. To this purpose we took advantage of the λN-boxB approach, where boxB sites were cloned upstream of the luciferase coding sequence and Mago was fused to λN. Upon expression of the fusion construct we found that Pol II exhibits significant increase in enrichment at promoter (*P* = 9.5 × 10^−5^) and a slight decrease at the 3’ end of luciferase, in comparison to expression of λN alone (Figure 2H). In contrast the expression of λN fused to GFP had no effect, while expression of λN-RnpS1 had an opposite effect regarding Pol II occupancy. Hence, these results indicate that recruiting Mago to the 5’ end of RNA is sufficient to increase promoter proximal pausing of RNA Pol II at the corresponding locus.

### 6 Loss of Mago Results in Global Changes in Chromatin Accessibility

Transcription is tightly coupled to chromatin architecture (Gilchrist et al., 2010). To address whether changes in Pol II pausing upon KD of pre-EJC components are associated with a change in chromatin organization, we examined the degree of chromatin compaction in wild type versus Mago-depleted cells. To this end, we performed MNase-Seq experiment, where chromatin was digested by micrococcal nuclease (MNase) and mono-nucleosomal fragments were subjected to paired-end deep sequencing. Examination of the chromatin structure at the genome wide level indicated extensive changes upon Mago depletion. The most notable change was at the level of TSS where nucleosomal occupancy was strongly increased (Figures 3A and 3B). This is consistent with previous findings indicating that paused Pol II and nucleosomes compete with each other at TSS (Gilchrist et al., 2010). Furthermore, the phasing of nucleosomes within the gene body was strongly altered (Figures 3A and 3B). Interestingly, pre-EJC-bound promoters show the most significant changes, as expected for a direct effect (Figure 3C, *P* < 2.2 × 10^−16^). A mild but significant negative correlation between changes in Ser2P level and chromatin accessibility was also found (Figure 3D, coefficient of determination R^2^ = −0.2274). Furthermore, we found reduced level of the activating histone mark H3K4me3 upon Mago KD, in particular at pre-EJC-bound genes (Figure 3E, *P* < 2.2 × 10^−16^). Altogether, these results indicate that Mago modulates histone marks and chromatin accessibility, likely as a consequence of its promoter proximal pausing activity.

**Figure 3.**
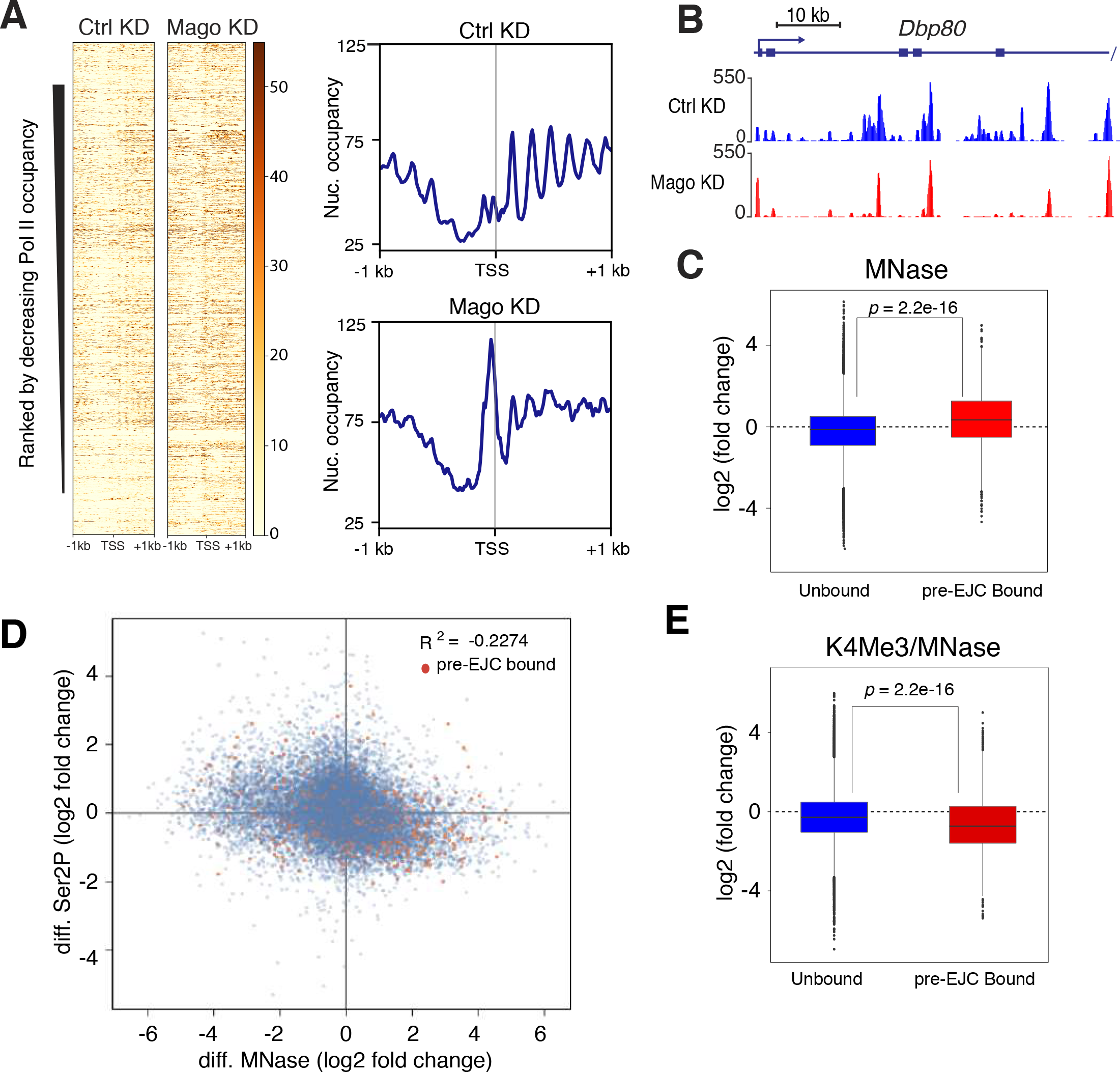
Mago Controls Chromatin Accessibility. **(A)** Heatmaps of nucleosomal occupancy from MNase-seq experiments performed in biological duplicates from S2R+ cells in control or Mago knockdown conditions, centered at the TSS in a ± 1 Kb window. Rows indicate all the genes bound by Pol II and are sorted by decreasing Pol II occupancy. The color labels to the right indicate the levels of nucleosomal occupancy. Composite metagene profiles are also shown, with nucleosomal occupancy level on Y-axis and genomic coordinates on X-axis. **(B)** Genome browser view of averaged MNase-seq data from two independent biological duplicates in S2R+ cells treated with control or Mago double stranded RNA. Example shown here is *Dbp80* gene, a well-defined pre-EJC target. **(C)** Log2 fold changes in nucleosomal occupancy at the TSS (250bp upstream and downstream of TSS) after control and Mago knockdowns. The changes were separated according to pre-EJC binding and a two-sample t-test was performed. **(D)** Scatterplot between changes in nucleosome and Ser2P occupancy after either control or Mago knockdown. Pre-EJC-bound promoters are highlighted by orange color. A mild negative correlation, as shown in the indicated pearson coefficient of correlation, between nucleosome and Ser2P occupancies was found. **(E)** Log2 fold changes in K4Me3 levels normalized to the nucleosomal occupancy (MNase data) at the TSS (250bp upstream and downstream of TSS), after control and Mago knockdowns. The changes were separated according to pre-EJC binding and a two-sample t-test was performed.

### 7 pre-EJC Gene Size Dependency is Mediated Transcriptionally

We showed that the expression of genes containing larger introns is more affected in the absence of pre-EJC components, while the loss of RnpS1 has no relationship with respect to intron size (Figure S1). We next examined whether underlying transcriptional changes might drive this size dependency. To address this, we classified genes into different intron classes as implemented previously, and calculated the fold changes in nucleosomal occupancy and Ser2P levels upon Mago depletion. These analyses revealed a striking pattern, whereby nucleosomal occupancy increases at promoters but decreases along the gene body and at the TES in an intron size dependent manner (Figure 4A). In contrast, Ser2P levels display anti-correlative changes with respect to the nucleosomal occupancy (Figure 4B), confirming our previous finding of a negative correlation between nucleosomal occupancy and Ser2P levels. We next analyzed changes in PRR upon Mago depletion using the same intron size classification. Strikingly, the changes in PRR also increase in a size dependent fashion (Figure 4C). Therefore, these results indicate that Mago has a stronger impact on the transcriptional regulation of genes with longer introns compared to the ones with shorter introns. In order to determine whether these changes in nucleosomal and Ser2P occupancy result from pre-EJC binding specificity, we calculated the percentage of genes bound by the pre-EJC in different classes relative to their representation in the total number of expressed genes. Interestingly, we found that pre-EJC binding was significantly over represented at genes containing longer intron (Figure 4D, *P* < 2.2 × 10^−16^). Consistent with a direct control of gene expression by pre-EJC components on long intron-containing genes, we found that expression of pre-EJC-bound genes was generally decreased upon KD of pre-EJC components (Figures 4E-G, *P* < 2.2 × 10^−16^) and this decrease was largely observed at nascent RNA (Figure 4H, *P* < 2.2 × 10^−16^). Collectively, our results indicate that pre-EJC components preferentially bind and regulate expression of large intron-containing genes via a direct transcriptional effect.

**Figure 4.**
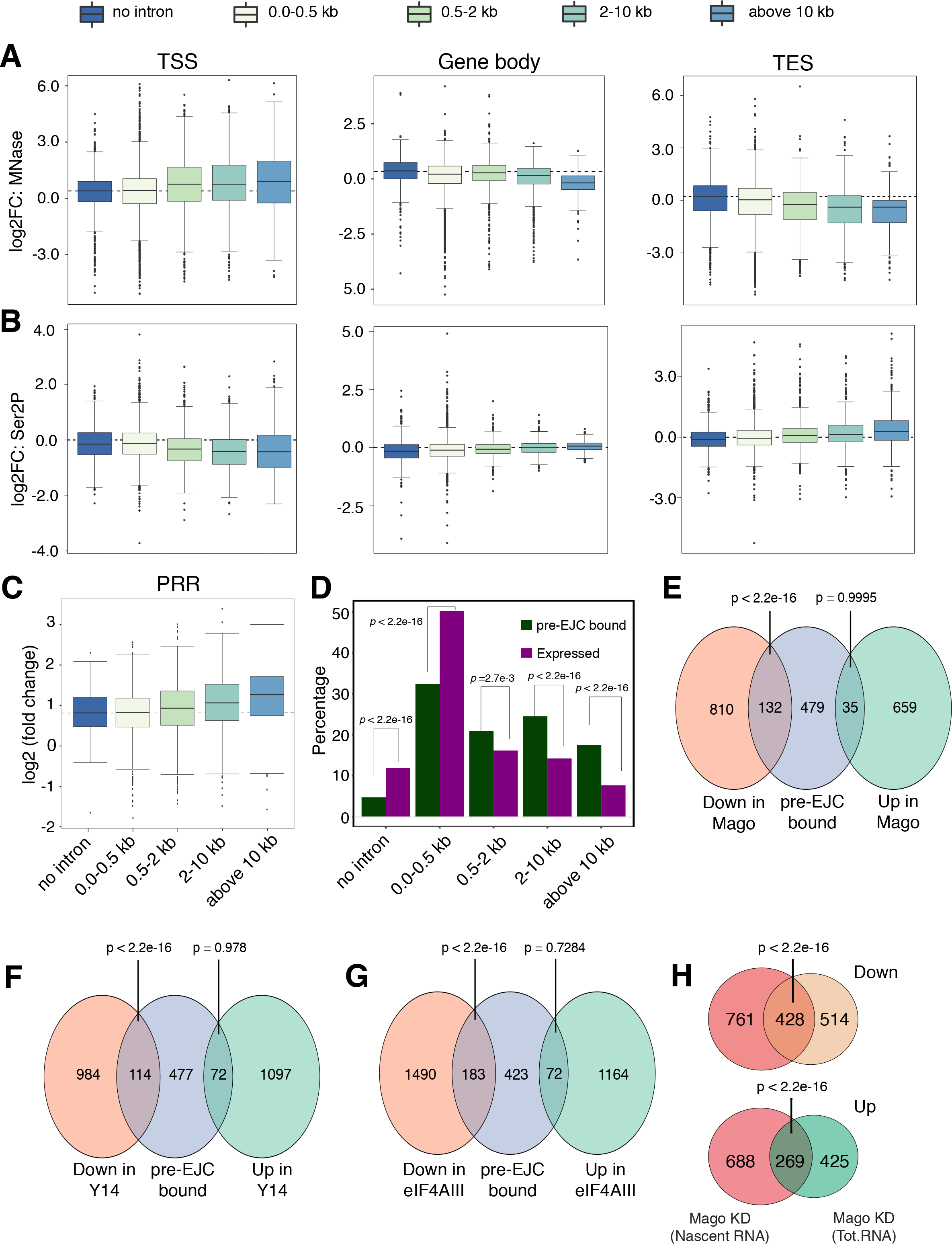
Mago Controls Chromatin Accessibility and Pol II Pausing in a Gene Size Dependent Manner. **(A, B)** Changes in nucleosome occupancy **(A)** and Ser2P levels **(B)** in control and Mago knockdown S2R+ cells at the promoter, gene body, and TES, separated according to the size of the largest intron (ANOVA, *p* < 2.2 × 10^−16^). **(C)** Box plots showing changes in PRRs upon Mago depletion when compared to control, separated into different intron classes. **(D)** Percentage of pre-EJC-bound genes in different intron classes, along with the percentage of each class amongst all the expressed genes. The proportion of pre-EJC-bound genes is increased for genes containing larger introns, relative to their abundance. *P*-values are derived from Fisher’s exact test. **(E, F, G)** Venn diagrams showing the overlap between Mago-, Y14- and eIF4A3-bound genes and genes with differential expression upon respective knockdowns. The overlap between genes that are down-regulated upon each pre-EJC component knockdown, and their respective target genes is significant. *P*-values are derived from Fisher’s exact test. **(H)** Venn diagram showing the overlap between genes identified as differentially expressed in mRNA sequencing and nascent RNA sequencing (4sU-Seq) upon Mago KD, separated for either up regulated and down regulated genes. *P*-values are derived from Fisher’s exact test.

### 8 Mago Stabilizes Promoter Proximal Pausing by Restricting P-TEFb Binding to Pol II

P-TEFb is a critical complex required to promote Ser2 phosphorylation and Pol II release. To determine whether the observed increase in Pol II release into the gene body results from changes in P-TEFb occupancy, we attempted to perform ChIP-Seq of Cdk9-HA-Flag in control cells or cells depleted of Mago. However, we observed only low Cdk9 enrichment on chromatin with poor reproducibility among different biological replicates (data not shown). This was likely due to weak association of Cdk9 with chromatin. To circumvent this issue, we took advantage of the targeted DamID (TaDa) approach to monitor Cdk9 occupancy. In contrast to the ChIP method DamID allows to capture weak and/or transient interactions (van Steensel and Henikoff, 2000; Vogel et al., 2007). Expression of N-terminally tagged Dam-Cdk9 was induced in S2R+ cells for 16 hours in control and Mago-depleted cells and DNA was further prepared and subjected to sequencing. Interestingly, we found a substantial increase in Cdk9 enrichment in Mago KD condition, compared to control KD (Figure 5A). Track examples of Cdk9-DamID normalized to DamID are shown in control and Mago KD conditions, along with Mago binding (Figure 5B). Furthermore, heatmap of Cdk9 enrichment revealed slightly higher occupancy upon Mago KD, when sorted according to decreasing Pol II occupancy (Figure 5C). Importantly, western blot analysis showed that the overall level of Cdk9 was unchanged upon Mago KD, indicating that this increase does not result from changes in protein expression (Figure S6A). Altogether, our results suggest that Mago controls Ser2 phosphorylation and Pol II pausing by restricting Cdk9 recruitment at promoter regions. To evaluate whether the change in Cdk9 occupancy was directly driven by Mago occupancy, we again divided promoter of expressed genes into pre-EJC-bound and unbound class, and plotted the change in Cdk9 enrichment for these two classes. The increase in Cdk9 enrichment for pre-EJC-bound class was mild albeit significantly higher than the unbound class (Figure 5D, *P* = 0.02508). Together, these results indicate that pre-EJC binding at promoter regions restricts Cdk9 occupancy at the targeted genes.

**Figure 5.**
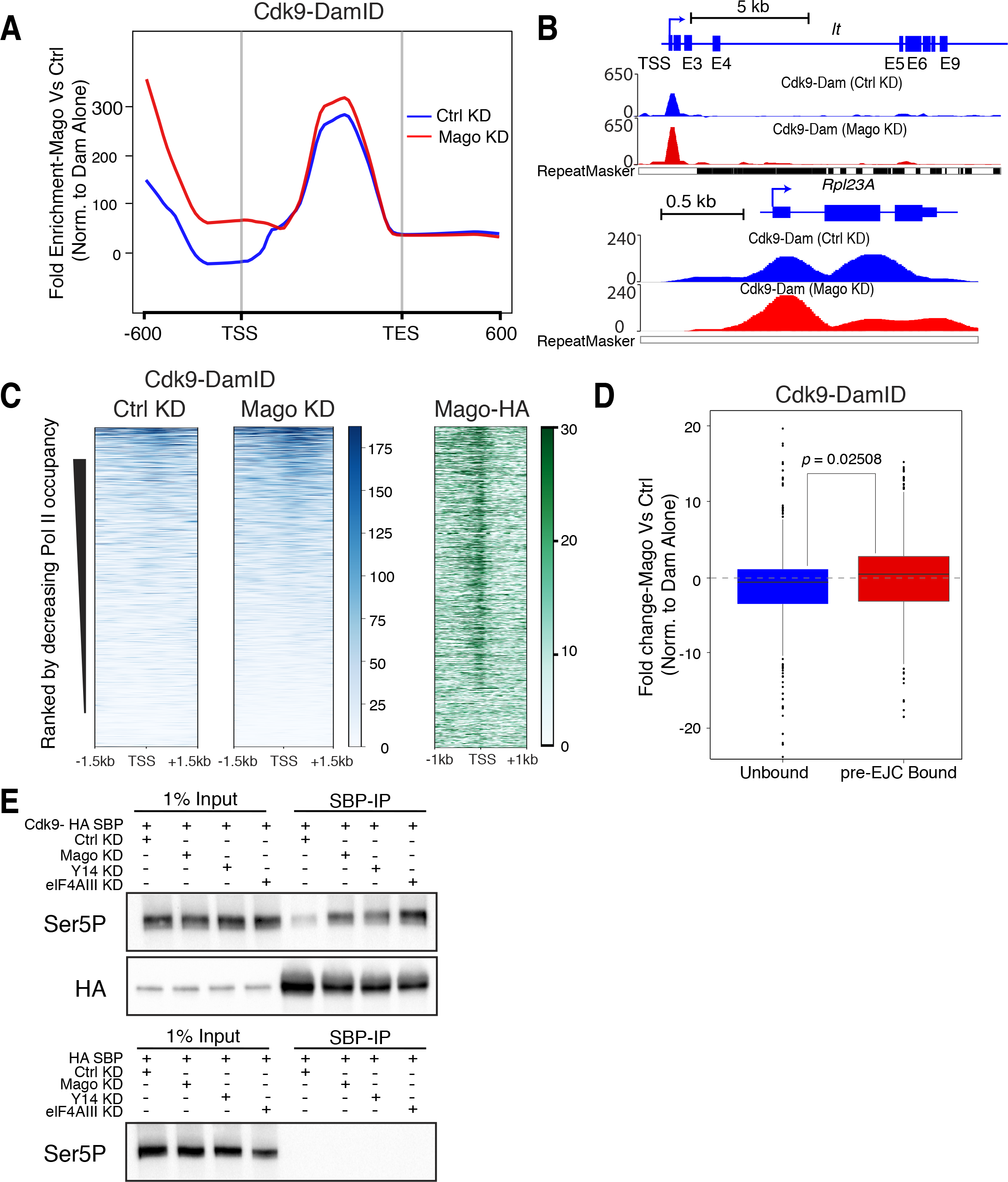
Loss of pre-EJC Core Components Results in Increased Cdk9 Binding to Pol II. **(A)** Averaged metagene profiles of Cdk9-DamID in control and Mago knockdown cells, from two independent biological replicates. Fold enrichment represented on Y-axis was calculated against Dam alone control in respective conditions (using damidseq_pipeline), while X-axis depicts scaled genomic coordinates. **(B)** Genome browser view of Dam alone normalized and averaged tracks of Cdk9-DamID for *light* and *Rpl23A*. **(C)** Heatmaps of normalized Cdk9-DamID enrichment, in either control or Mago depleted S2R+ cells, centered at the TSS in a ± 1.5 Kb window. Rows indicate all the genes bound by Pol II and are sorted by decreasing Pol II occupancy, and the color labels to the right indicate the level of enrichment. Heatmap of Mago-HA centered at the TSS (−1kb to +1kb) is also shown. **(D)** Changes in Cdk9 occupancy at the TSS (250bp upstream and downstream of TSS) after control and Mago knockdowns, calculated from Cdk9-DamID experiment, after normalizing to the Dam alone control. The changes were separated according to pre-EJC binding and a two-sample t-test was performed. **(E)** Co-immunoprecipitation experiments from S2R+ cells extracts expressing either HA-SBP-Cdk9 or HA-SBP alone, revealed with Pol II Ser5P antibody. Immunoprecipitations were performed from control cells or cells depleted for pre-EJC core components (Mago, Y14, and elF4AIII). Shown also is the western blot against HA tag for assessing the efficiency of the pull-down of HA-SBP-Cdk9 in each knockdown condition.

To address whether Mago might restrict P-TEFb binding to Pol II, we immunoprecipitated Cdk9 with a HA-SBP tag in either control condition or upon Mago depletion, and tested for the presence of the Ser5 phosphorylated form of Pol II by western blot analysis. Notably, a substantial increase of Cdk9 binding to Pol II Ser5P was observed when Mago levels were reduced (Figure 5E). This increase was not due to changes in immunoprecipitation efficiency, as Cdk9 was similarly precipitated in control versus Mago KD conditions. We found similar results when other pre-EJC components were depleted, strongly suggesting that the pre-EJC restricts binding of P-TEFb to Pol II, which in turn reduces Ser2P levels and the entry of Pol II into elongation.

### 9 Reducing Pol II Release is Sufficient to Rescue Mago Defects *in vivo*

Accumulating evidences indicate that promoter proximal pausing of Pol II may serve as a checkpoint to influence downstream RNA processing events (Adelman and Lis, 2012; Barboric et al., 2009; Ghosh et al., 2011; Mandal et al., 2004; Moore and Proudfoot, 2009; Rasmussen and Lis, 1993). Therefore, to test whether the decrease of Pol II pausing accounts for some of the exon skipping observed upon Mago KD we attempted to rescue the splicing defects by decreasing the release of Pol II into gene bodies. To this end, we simultaneously depleted Cdk9 and Mago in S2R+ cells and analyze the effect on pre-mRNA splicing. We first noticed that Cdk9 KD rescues the increase of Ser2P levels observed upon Mago-depletion by western blotting (Figure 6A). To further evaluate the extent of the rescue at the gene level, we performed ChIP-qPCR using Ser2P antibody and examined the *MAPK* locus. This experiment revealed a partial rescue of Ser2P occupancy at the *MAPK* gene (Figure 6B), indicating that Cdk9 counteracts the effect of Mago in controlling Ser2P levels. Consistent with this counteracting role at the transcriptional level, the gene size-dependency relative to expression levels observed upon Mago KD was lost in the double KD (Fig. S6B).

**Figure 6.**
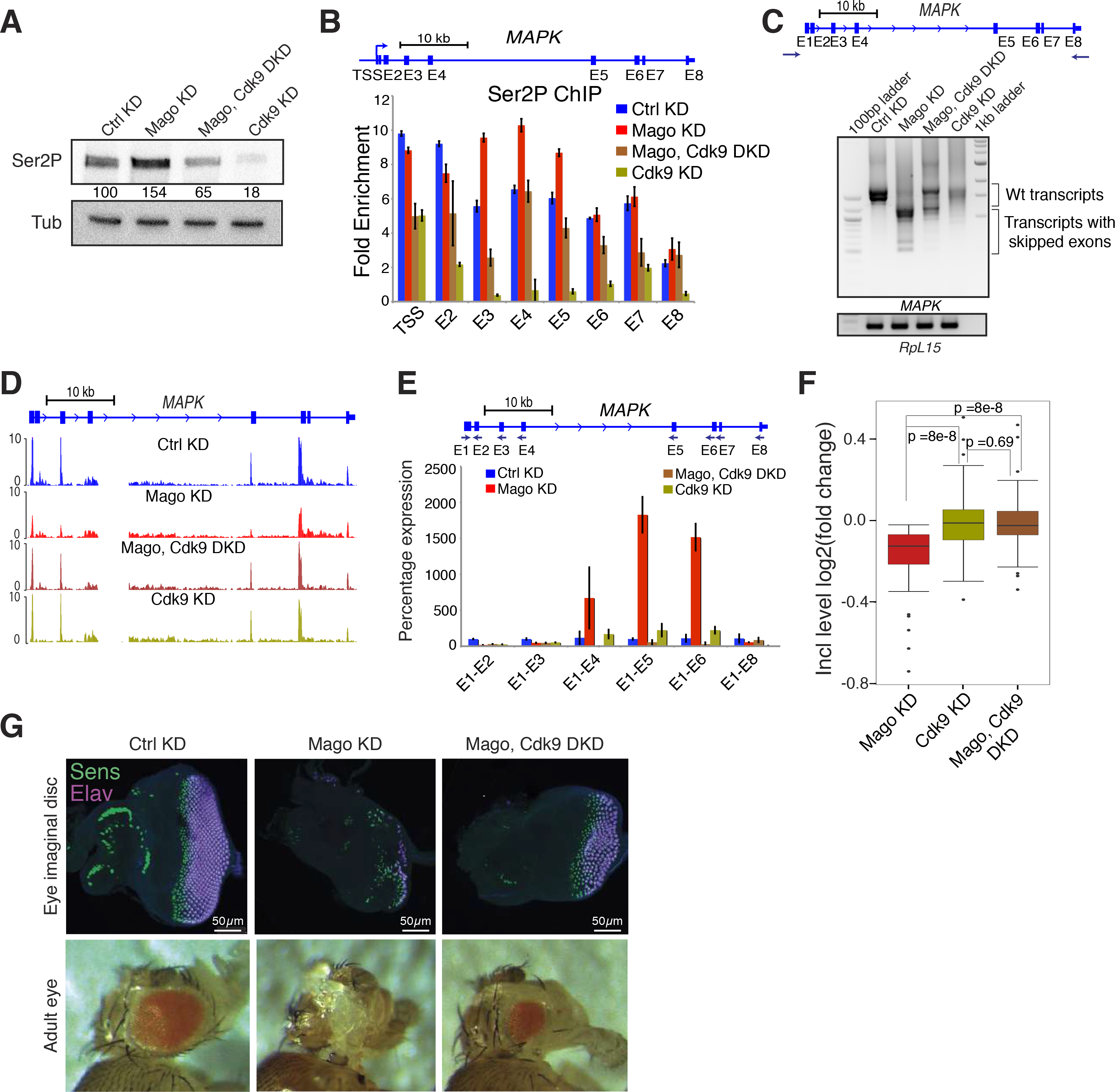
Restoring Pausing is Sufficient to Rescue Mago-Associated Exon Skipping Defects. **(A)** Mago knockdown results in elevated level of Ser2P phosphorylation of Pol II. Western blotting using antibodies against Pol II Ser2P and Tubulin, using S2R+ cell extracts with indicated knockdowns. Signal in the knockdown conditions was normalized to the control condition using Tubulin as loading control and quantification of the intensity was performed with ImageJ. **(B)** ChIP-qPCR analysis of Ser2P occupancy level at *MAPK* locus in the indicated knockdowns. The primers used for the analysis are indicated in the scheme above. Error bars indicate the standard deviation from the mean of two independent biological replicates. **(C)** Agarose gels showing RT-PCR products for *MAPK* using RNA extracted from S2R+ cells with indicated knockdowns as template for cDNA synthesis. The primers used for the PCR 5’ and 3’ UTR of *MAPK*, as shown in the scheme above. RT-PCR products for *RpL15* from respective knockdown condition were used as loading control. **(D)** Replicate averaged RNA-Seq track examples of *MAPK* in several knockdown conditions. Mago KD results in several exon skipping events, which are rescued upon simultaneous knockdown of Cdk9. **(E)** Quantitative RT-PCR using RNA extracts derived from S2R+ cells with the indicated knockdowns. The amplicons were obtained using the same 5’ forward primer (E1) together with the reverse primers on respective exons, as shown in the scheme above. The level of exon skipping is compared to the control treatment, with *RpL49* used for normalization. Error bars indicate the standard deviation from the mean of two biological replicates. **(F)** Box plots showing the log2 fold change in inclusion level of alternatively spliced exons in the indicated knockdowns (rMATS was used for the analysis). **(G)** Upper panel: Staining of eye imaginal discs from third instar larvae with indicated dsRNAs specifically expressed in the eye (using the *ey*-GAL4 driver). All photoreceptors are stained with anti-Elav antibody (purple), and R8 (the first class of photoreceptor to be specified) with anti-Senseless (green). Lower panel: Adult eyes of a control fly and flies with indicated KD.

We next explore the effect of depleting Cdk9 on Mago-mediated splicing events. Strikingly, reducing Cdk9 levels almost fully rescued *MAPK* splicing, as assessed by semi-quantitative RT-PCR (Figure 6C). This was confirmed by RNA-Seq and quantitative RT-PCR experiments (Figures 6D and 6E). Remarkably, this rescuing effect was not only restricted to *MAPK* but was also observed for other Mago-dependent exon skipping events (Figure 6F, *P* =8 × 10^−8^). Consistent with an effect of Mago on splicing via modulation of promoter proximal pausing we found that genes that display differential splicing upon Mago KD were significantly enriched for pre-EJC binding (Figure S6C, Fisher’s test p < 2.2 × 10^−16^). We could extend this observation to other pre-EJC components (Figures S6D and S6E). Additionally, we found that depleting Cdk12, another kinase recently involved in the release of promoter proximal pausing (Yu et al., 2015), rescued *MAPK* splicing of Mago-depleted cells to a similar extent as with Cdk9 KD (Figures S6F and S6G). Altogether, these results strongly suggest that Mago regulates gene expression and exon definition via regulation of Pol II promoter proximal pausing.

Given that *MAPK* is the main target of the EJC during *Drosophila* eye development (Roignant and Treisman, 2010), we attempted to rescue the photoreceptor differentiation defect of Mago-depleted cells by simultaneously reducing Cdk9 levels specifically in the eye. As shown previously, eye-specific depletion of Mago strongly impairs photoreceptor differentiation (Ashton-Beaucage et al., 2010; Roignant and Treisman, 2010). However, decreasing Cdk9 function in a similar background remarkably rescued eye development (Figure 6G). Notably, the number of differentiated photoreceptors in larvae and adults was substantially increased. Similar rescue of photoreceptor differentiation was observed in the double KD for Mago and Cdk12 (Figure S7A). This effect was not restricted to Mago as depletion of Cdk9 was also sufficient to rescue the lethality and the eye defects associated with eIF4AIII KD (Figure S7B). However, reducing the speed of Pol II or depleting several transcription elongation factors (lilli, enl, ell, atms, cdc73, SSRP1, dre4 and rtf1) failed to substantially rescue eye development in absence of Mago (Figures S7C-S7F), providing additional evidence that Mago transcriptional function occurs at the level of promoter proximal pausing rather than at the transcription elongation stage. Altogether, these results demonstrate that - despite the numerous post-transcriptional functions of the EJC - simply modulating Pol II release is sufficient to rescue the eye defect associated with Mago depletion.

### 10 The Function of Mago in Promoter Proximal Pausing is Evolutionary Conserved

To check whether EJC-mediated promoter proximal pausing is conserved in vertebrates we performed ChIP-Seq in HeLa cells for Pol II upon depletion of Magoh, the human ortholog of *Drosophila* Mago. Similar to our observation in S2R+ cells, we found that decreased levels of Magoh led to an increased release of Pol II from promoter to the gene body that translates into higher PRR (Figures 7A-C). The Track examples illustrate the lower Pol II enrichment at the promoter and higher in the genebody, in Magoh knockdown compared to control condition (Figure 7D). Consistent with this observation and with our findings in S2R+ cells, western blot analysis revealed higher level of Ser2P while Ser5P remains unaffected (Figure 7E). We also find that Magoh specifically interacts with Pol II and with Ser5P but fails to associate with Ser2P (Figures 7F-H). We further tested whether Magoh depletion similarly affects the interaction between the Ser5 phosphorylated form of Pol II and Cdk9. We performed Cdk9 co-immunoprecipitation in control and Magoh depleted conditions and tested for the presence of Ser5P. Consistent with our findings in S2R+ cells, a stronger interaction between Ser5P and Cdk9 was observed upon Magoh KD (Figure 7I). Thus, these data indicate that the function and mechanism of the pre-EJC in the control of promoter proximal pausing is conserved in human cells.

**Figure 7.**
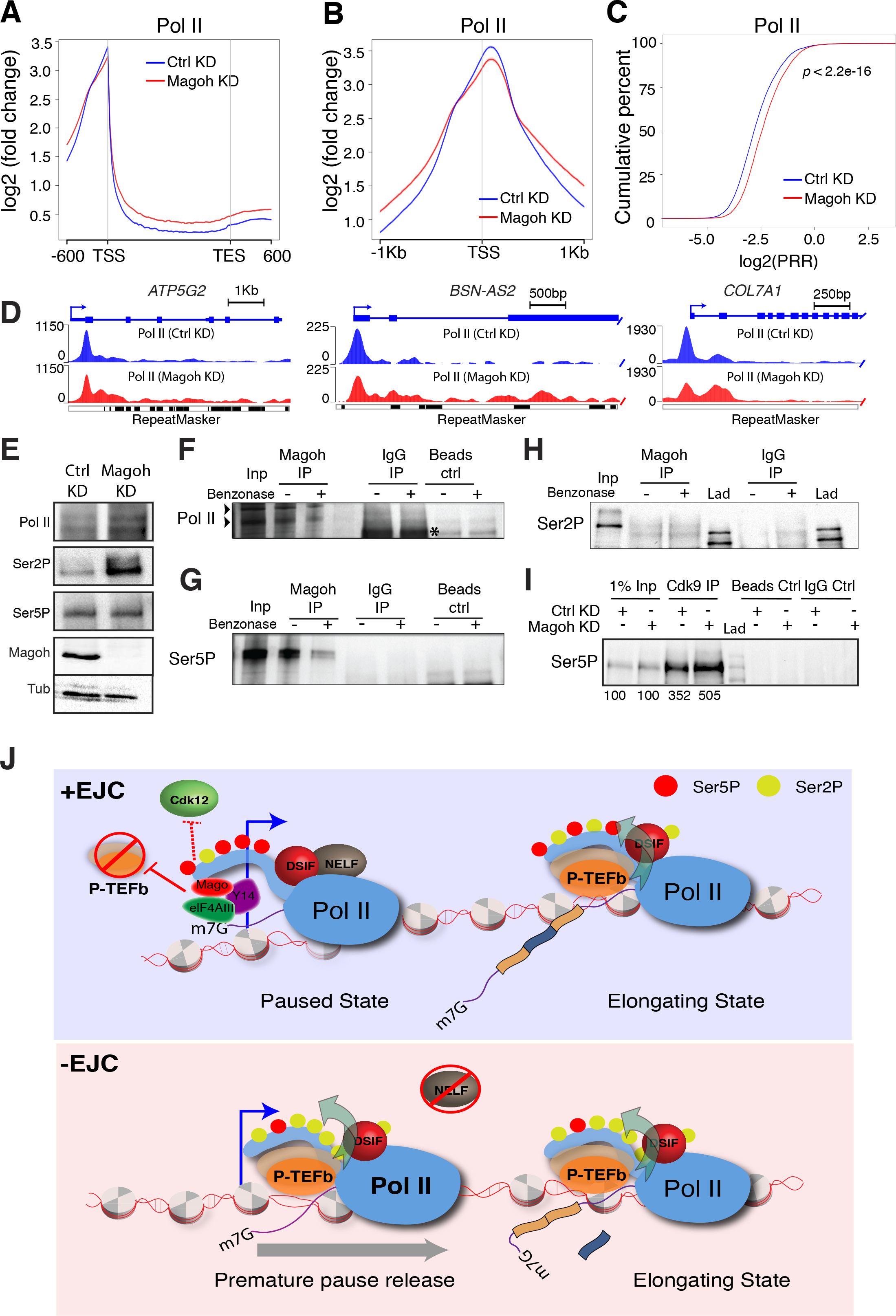
The role of Mago on Promoter Proximal Pausing is Conserved in Human. **(A)** Averaged metagene profiles from two independent biological replicates of total Pol II occupancies in control and Magoh-depleted HeLa cells. Log2 fold changes against input control are shown on Y-axis while X-axis depicts scaled genomic coordinates. **(B)** Averaged metagene profiles from two independent biological replicates of total Pol II occupancies in control and Magoh-depleted HeLa cells, centered at the TSS in a ± 1 Kb window. Log2 fold changes against input control are shown on Y-axis while X-axis depicts genomic coordinates. **(C)** The ECDF plot of PRR in HeLa cells treated with either control or *Magoh* siRNA. *P*-value is derived from two-sample Kolmogorov-Smirnov test. **(D)** Track examples of total Pol II ChIP-Seq from HeLa cells extracts, after either control or Magoh knockdown. The tracks are average of two independent biological replicates after input normalization. **(E)** Western blotting performed using antibodies against total Pol II, Ser2P, Ser5P and Magoh, in HeLa cells treated with either control or *Magoh* siRNA. Similar to *Drosophila*, the loss of Magoh leads to elevated level of Ser2P phosphorylation without affecting Ser5P phosphorylation. **(F, G, H)** Co-immunoprecipitation of Magoh from HeLa cells extracts, using antibody directed against Magoh. Western blots using Pol II, Ser5P and Ser2P antibody reveal RNA-dependent interaction of Magoh with Pol II and Ser5P. There was no detectable interaction of Magoh with Ser2P phosphorylated form of Pol II. The specific bands for Pol II are highlighted by arrowheads in F. The lane labeled with “Lad” indicates ladder. **(I)** Co-immunoprecipitation of Cdk9 using anti-Cdk9 antibody from HeLa cells extracts treated with either control or *Magoh* siRNA. Western blot was performed with anti-Ser5P antibody. The quantification of the intensity was performed with ImageJ **(J)** Model: The EJC stabilizes Pol II pausing by restricting P-TEFb binding at promoter, and possibly by sequestering Cdk12. This activity is required for proper recognition of exons.

## DISCUSSION

Our work uncovers an unexpected connection between the nuclear EJC and the transcription machinery via the regulation of Pol II pausing, which is conserved from flies to human. The pre-EJC stabilizes Pol II in a paused state at least in part by restricting the association of P-TEFb with Pol II via non-canonical binding to promoter regions. The premature release of Pol II into elongation in absence of the EJC results in splicing defects, highlighting the importance of this regulatory step in controlling downstream RNA processing events (Figure 7J).

### The EJC Stabilizes Pol II in a Paused Configuration

Promoter proximal pausing is a widespread transcriptional checkpoint, whose functions and mechanisms have been extensively studied. Several important regulators have been identified, which includes the positive transcription factor P-TEFb and the negative regulators NELF and DSIF. Our data reveal that the pre-EJC plays similar roles as the previously described negative factors by preventing premature pol II release into elongation. How does the pre-EJC control pol II pausing and how does it interplay with other pausing regulators? Our study provides some answers to these questions. In absence of pre-EJC components, P-TEFb associates more strongly with Pol II, which results in increased Ser2 phosphorylation, demonstrating that one of the activities of the pre-EJC is to restrain P-TEFb function by diminishing its association with chromatin. While it is not clear yet how the pre-EJC exerts this function, a simple mechanism would be by steric interference for Pol II binding, although more indirect mechanisms might also exist. This mechanism infers that both the pre-EJC and Cdk9 bind similar sites on the CTD on Pol II, which fits with the association of the pre-EJC with the Ser5 phosphorylated form of pol II and not with Ser2P. However, we also found elevated Cdk9 binding and premature release of Pol II at Mago-unbound genes, albeit to a lesser extent compared to Mago-bound genes, suggesting that additional mechanisms must be involved. The fact that the KD of Cdk12 also rescues Mago splicing defects supports this possibility.

It is interesting to note that the binding of the pre-EJC to Pol II requires the presence of nascent RNA. A recent study also supports these findings showing specific association of pre-EJC components on polytene chromosomes that depends on nascent transcription but is independent of splicing (Choudhury et al., 2016). This is reminiscent to the binding of DSIF and NELF (Battaglia et al., 2017; Blythe et al., 2016; Cheng and Price, 2008; Crickard et al., 2016; Lee et al., 2008; Narita et al., 2003), suggesting that interaction with pol II and stabilization via nascent RNA is a general mechanism to ensure that pausing regulators exert their function at the right time and at the right location. Upon external cues, P-TEFb modifies the activities of both NELF and DSIF through phosphorylation, promoting pol II release. It would be of interest to address whether P-TEFb also regulates the EJC in a similar manner. Intriguingly, previous studies revealed that eIF4AIII is present in the nuclear cap-binding complex (Choe et al., 2014), while Y14 directly recognizes and binds the mRNA cap structure (Chuang et al., 2013; Chuang et al., 2016). It is therefore possible that this cap-binding activity confers the ability of the EJC to bind nascent RNA. Nevertheless, other factors must be clearly involved as only a subset of genes is bound by the EJC. We speculate that previous *in vitro* transcription experiments aimed at identifying novel pausing factors might have failed to detect pre-EJC binding at promoter regions due to its very weak association to Pol II in the absence of RNA. Hence our results highlight the need to embark into more *in vivo* genetic studies to identify additional regulators of this major transcription regulatory step.

SRSF2 is another splicing regulator that was previously demonstrated to modulate pol II pausing via binding to nascent RNAs (Ji et al., 2013). In this case, SRSF2 exerts an opposite effect by facilitating pol II release into the elongation phase. This effect occurs via increased P-TEFb recruitment to gene promoters. Although we have not found convincing evidence for a conserved role of the *Drosophila* SRSF2 homolog in this process (unpublished data), one may envision that the pre-EJC counteracts the effect of SRSF2 to stabilize pol II pausing. Consistent with this possibility, EJC binding sites are often associated with RNA motifs that resemble the binding sites for SR proteins (Singh et al., 2012). It is therefore possible that SR proteins influence pre-EJC loading to mRNA and vice versa. More experiments will be required to address the link between the pre-EJC and SR proteins in the regulation of promoter proximal pausing.

### Role of the EJC in Alternative Splicing

Our previous work along with studies from other groups suggested that the EJC modulates splicing by two distinct mechanisms (Ashton-Beaucage et al., 2010; Hayashi et al., 2014; Malone et al., 2014; Roignant and Treisman, 2010). On one hand, the EJC facilitates the recognition and removal of weak introns after prior deposition to flanking exon-exon boundaries. We proposed that EJC deposition occurs in a splicing dependent manner after rapid removal of *bona fide* introns, which are present in the same transcript. Thus, a mixture of “strong” and “weak” introns ensures EJC’s requirement in helping intron definition. This function requires the activity of the EJC splicing subunits Acinus and RnpS1, which are likely involved in the subsequent recruitment of the splicing machinery near the weak introns. While this model is attractive, it does not however explain every EJC-regulated splicing event. In particular, depletion of pre-EJC components results in a myriad of exon-skipping events, which occur frequently on large intron-containing transcripts (this study and (Ashton-Beaucage et al., 2010; Roignant and Treisman, 2010)). In contrast to intron definition, this exon definition activity only slightly required the EJC splicing subunits, suggesting an additional mechanism. We now show that the pre-EJC controls exon definition by preventing premature release of Pol II into transcription elongation. Our results shed light on a recent observation in human cells showing that the usage of general transcription inhibitors improve splicing efficiency on two EJC-mediated exon skipping events (Wang et al., 2014).

The notion that splicing takes place co-transcriptionally is now a general consensus and two non-exclusive models regarding the impact of transcription, in particular of Pol II, on splicing have been proposed. Through the ability of the C-terminal repeat domain (CTD) of its large subunit, RNA Pol II can recruit a wide range of proteins, including splicing factors, to nascent transcripts (Buratowski, 2009; de Almeida and Carmo-Fonseca, 2008; Misteli and Spector, 1999; Phatnani and Greenleaf, 2006), thereby influencing intron removal. Pol II can influence splicing via a second mechanism referred to as kinetic coupling. According to the model, decreased elongation rates enhance the recognition of exons containing weak splice sites that would otherwise be skipped by the splicing machinery (Kornblihtt, 2006, 2007). This model was recently refined, as such that both increase and decrease of elongation rate can lead to splicing defects (Fong et al., 2014). In regards to pre-EJC’s activity we favor the first model. First, our genome wide studies demonstrate a global impact of the pre-EJC on promoter proximal pausing. Second, despite the premature release of Pol II into elongation, we did not observe substantial alteration of the average rate of transcription elongation upon Mago depletion. We did find however some gene-to-gene differences but they poorly correlate with the degree of exon inclusion. Still, this effect might be a secondary consequence of splicing defects, as a previous study suggested the existence of splicing-dependent elongation checkpoint (Chathoth et al., 2014). Third, in contrast to the depletion of P-TEFb, reducing the speed of Pol II or depleting the function of transcription elongation factors failed to rescue Mago-splicing defects, arguing that the positive impact of reducing P-TEFb levels on exon definition is dependent on its function in Pol II release rather than in regulating the elongation stage. Thus, in the light of previous model regarding the interplay between pre-mRNA capping and transcription, we propose that by stabilizing Pol II pausing the EJC provides enough time for the recruitment of additional splicing factors that play critical role in exon definition.

Pol II pausing is found more prominently at developmentally regulated genes, which tend to be long and frequently regulated by alternative splicing. We found a size dependency for Mago-bound genes as well as for Mago-regulated gene expression, suggesting that pre-EJC function is adapted to regulate exon definition of large genes by enhancing their promoter proximal pausing. Interestingly, a recent study shows that genes with long introns tend to be spliced faster and more accurately (Pai et al., 2017). Whether this function depends on EJC binding to nascent RNA constitutes an interesting possibility. The next important challenge will be to address the precise mechanisms by which promoter proximal pausing influences pre-mRNA splicing at these developmental genes.

## ACKNOWLEDGMENTS

We thank the Bloomington Drosophila Stock Center, the Transgenic RNAi Project in Harvard and the Vienna Drosophila Resource Center for fly reagents; the IMB Genomics and Bioinformatics Core facilities for tremendous support; members of Ulrich lab, especially Lilliana Batista for help with yeast chromatin for “spike-in” control; members of the Roignant lab for fruitful discussion; and Enrico Cannavo, Yad Ghavi-Helm, Guillaume Junion, Jessica Treisman for critical reading of the manuscript. This work was supported by the Marie Curie CIG 334288.

## AUTHOR CONTRIBUTIONS

J.A and J.Y.R designed the experiments. J.A performed the experiments and analyzed the data, D.B helped with RIP experiments. N.K, F. M., J.A and H.B performed the bioinformatics and statistical analyses. G.M.K carried out the *in vivo* and cell culture rescue experiments. J.A and J.Y.R wrote the paper.

## SUPPLEMENTARY FIGURE LEGENDS

**Figure S1.**
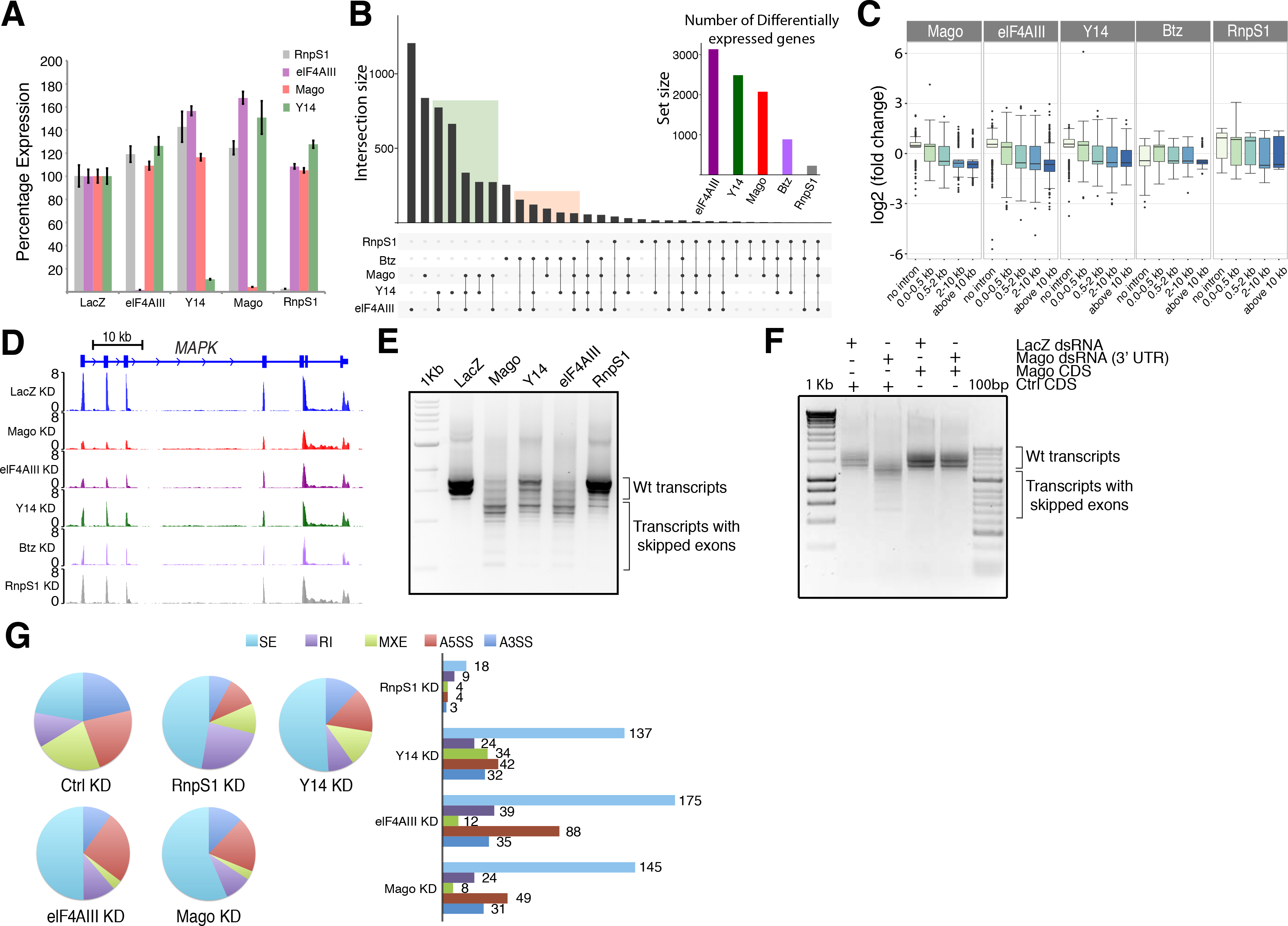
The pre-EJC affects expression and splicing of large intron-containing transcripts, related to Figure 1. **(A)** RT-qPCR analysis showing the efficiency of knockdown for the indicated conditions. **(B)** UpSet plot showing the overlap between differentially expressed genes in indicated knock down conditions. The Y-axis shows the size of the intersection while the X-axis shows different conditions in which the overlap was observed, highlighted by line connecting the solid dots. The histogram in the inset depicts number of differentially expressed genes in each condition. The overlap between at least two pre-EJC and two EJC components are highlighted with rectangular color shade. **(C)** Log2 fold changes in steady state RNA levels in indicated knockdowns compared to control knockdown, when genes are separated according to the size of their largest introns. The Y-axis shows the fold changes in expression while the X-axis depicts different classes **(D)**. Genome browser view of averaged steady state RNA-Seq data for *MAPK* transcript from S2R+ cells in the indicated knockdowns. Reads per million are shown on Y-axis. **(E)** Agarose gel of semi-quantitative RT-PCR for *MAPK* transcripts using RNA from S2R+ cells in the indicated knockdowns. Note that the depletion of pre-EJC core components (Mago, Y14, and elF4AIII) has greater effect on exon definition than RnpS1. **(F)** Agarose gel of semi-quantitative RT-PCR for *MAPK* transcripts using RNA from S2R+ cells with dsRNA targeting the 3’ UTR of Mago. The knockdown was performed in S2R+ cells either transfected with a control CDS or Mago CDS without 3’ UTR. Note that knockdown in cells transfected with Mago CDS does not lead to splicing defects in *MAPK*. The primers used for the PCR anneals in the 5’ and 3’ UTR of *MAPK*, as described before. **(G)** rMATS analysis of steady state RNA-Seq from S2R+ cells in the indicated knockdowns. (Left) The pie-charts depicts the percentage of significant events with skipped exons (SE), retained introns (RI), mutually exclusive exons (MXE), alternative 5’ splice site (A5SS) and alternative 3’ splice site (A3SS) in each condition, along the control with proportion of all events. (Right) Histograms showing the number of these events in indicated knockdowns when compared to control.

**Figure S2.**
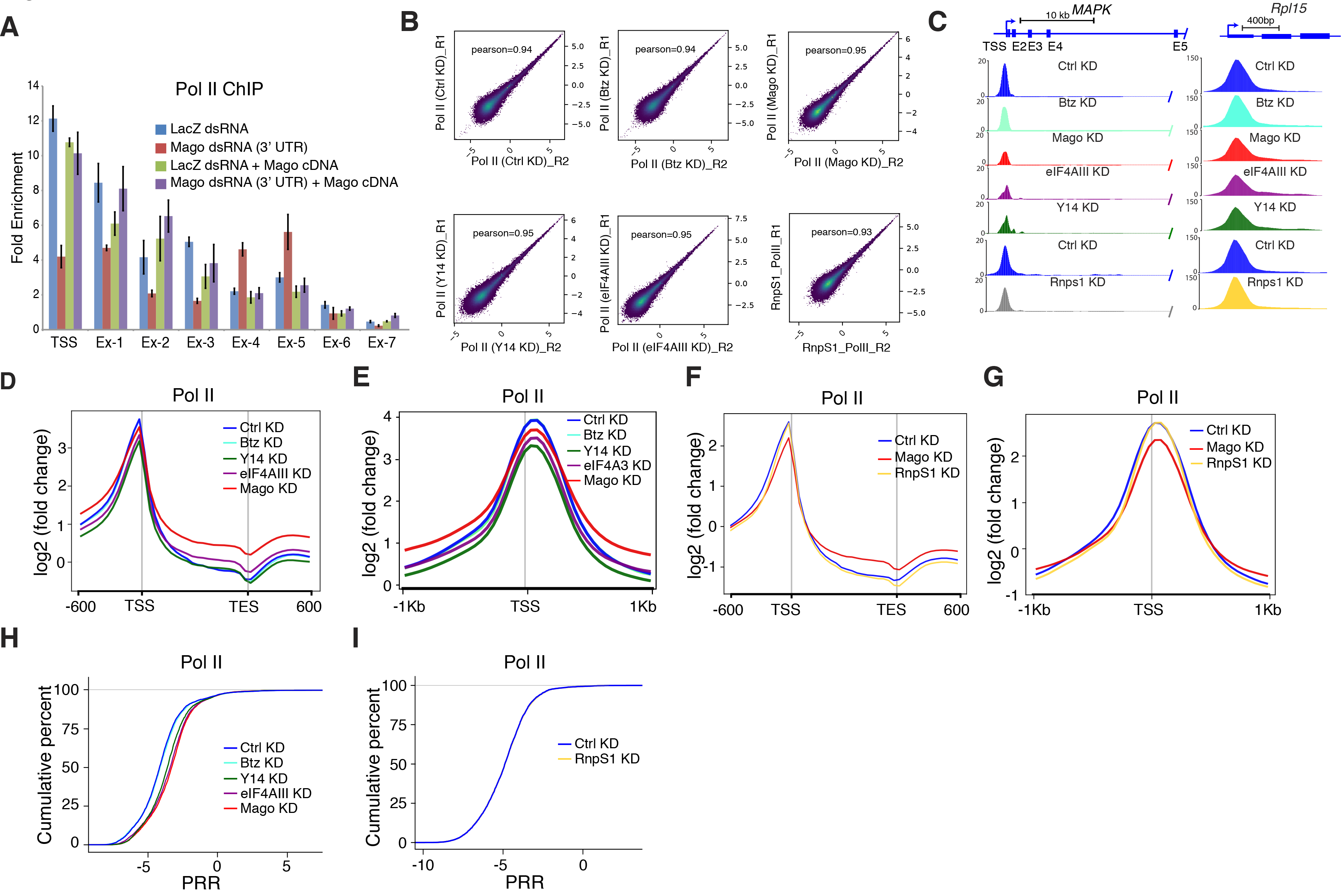
The EJC Modulates Promoter Proximal Pausing Independently of Btz and of its Splicing Subunit RnpS1, related to Figure 1. **(A)** ChIP-qPCR analysis of Pol II occupancies at *MAPK* locus. The tested regions for enrichment are same as shown in figure 1A. The knockdown was performed in S2R+ cells with dsRNA targeting the 3’ UTR of Mago, either transfected with a control CDS or Mago CDS without 3’ UTR. Note that knockdown in the cells transfected with Mago CDS does not lead to change in Pol II occupancies. Error bars indicate the standard deviation from the mean in three biological replicates. **(B)** Pearson correlation between the replicates of Pol II ChIP-Seq for indicated knockdowns. **(C)** Input and “spike-in” normalized and average track examples of total Pol II ChIP-Seq from S2R+ cells extracts, after either control or indicated knockdowns. Shown here are *MAPK* and Rpl15 loci. **(D)** Averaged metagene profiles from two independent biological replicates of total Pol II occupancies with “spike-in” normalization in control and indicated knockdown conditions. Log2 fold changes against input control are shown on Y-axis, while X-axis depicts scaled genomic coordinates. **(E)** Replicate averaged metagene profiles of total Pol II occupancies in control and in indicated knockdown conditions, after ““spike-in”” normalization, centered at the TSS in a ± 1 Kb window. Log2 fold changes against input control are shown on Y-axis, while X-axis shows genomic coordinates. **(F)** Averaged metagene profiles from two independent biological replicates of total Pol II occupancies with “spike-in” normalization in control, Mago and RnpS1 knockdown conditions. Log2 fold changes against input control are shown on Y-axis, while X-axis depicts scaled genomic coordinates. **(G)** Averaged metagene profiles from two independent biological replicates of total Pol II occupancies in control, Mago and RnpS1 knockdowns, after “spike-in” normalization, centered at the TSS in a ± 1 Kb window. Log2 fold changes against input control are shown on Y-axis, while X-axis shows genomic coordinates. **(H)** ECDF plots of PRR in cells with control and indicated knockdowns. KD of pre-EJC components has substantial effect on PRR while KD of the cytoplasmic component Btz has no effect. **(I)** ECDF plots of PRR in cells with control and RnpS1 knockdowns. The RnpS1 knockdown has no effect on the PRR.

**Figure S3.**
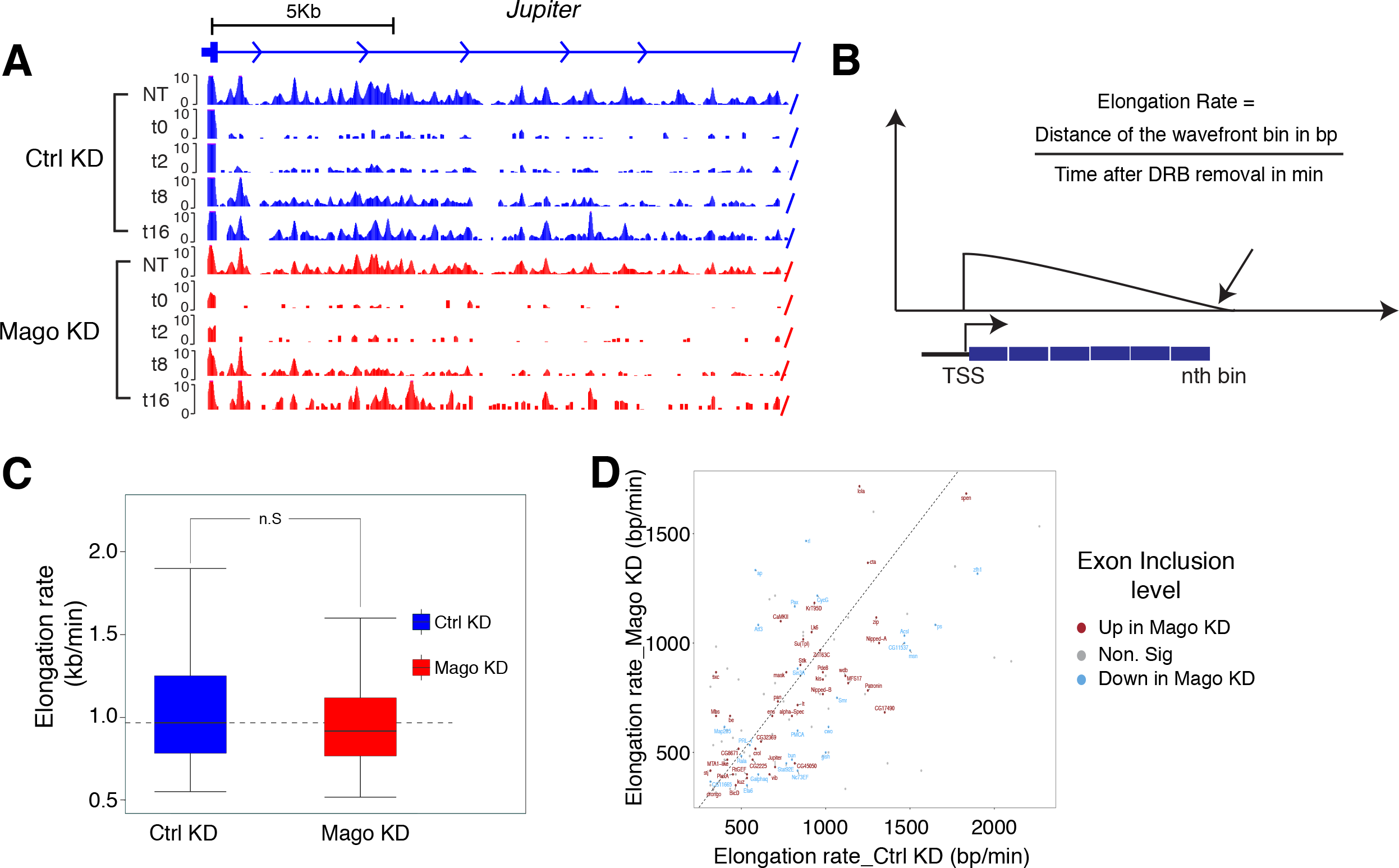
Mago Does Not Influence the Elongation Rate, related to Figure 1. **(A)** Genome browser view of DRB-4sU-Seq in control or Mago-depleted cells, shown here is *jupiter*. Time points indicate the time (in minutes) for which transcription was allowed to continue after removal of DRB. **(B)** Schematic representation of the calculation of the elongation rate. Genes longer than 10 kb were divided into 100 bp bins and the transcriptional wave front was identified in the bin with lowest local minima signal. The distance covered by the wavefront between 2 min after DRB removal and 8 min is then divided by the corresponding time interval (8 - 2 min) to calculate elongation rates. **(C)** Box plots showing the distribution of elongation rate in control and Mago-depleted S2R+ cells. **(D)** Scatterplot showing the relationship between elongation rate and exon inclusion level. The exon of the genes with the highest difference in inclusion level between control and Mago KD was considered for analysis (DEXseq was used for inclusion level estimates). The red and blue dots depict the genes with inclusion level in Mago knockdown higher and lower respectively, when compared to control knockdown. The grey dot depicts all those genes where the difference in inclusion level between the two conditions was not significant.

**Figure S4.**
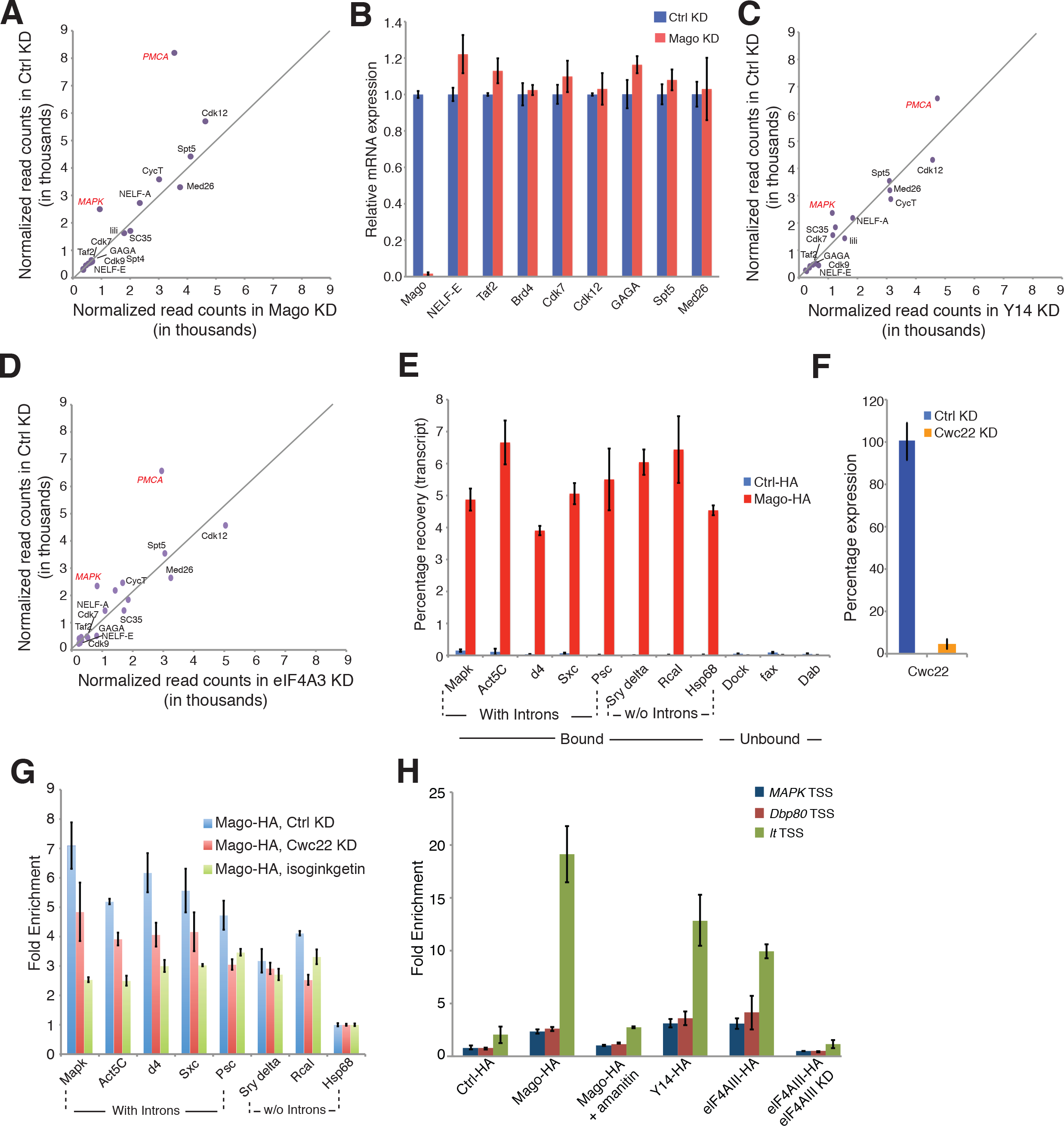
Pre-EJC Components Do Not Control the Expression of Paused and Elongation Factors, but Bind Nascent RNA, related to Figure 2. **(A)** Scatterplot showing the normalized read counts from DESeq2 of well-characterized elongation and paused factors in control and Mago-depleted S2R+ cells. Two Mago targets (*MAPK* and *PMCA*) are shown in red as controls. **(B)** Quantitative RT-PCR showing the transcript levels of genes involved in elongation and pause release control, using RNA extracts derived from S2R+ cells upon control and Mago knockdowns. Error bars indicate the standard deviation from the mean of two biological replicates. **(C, D)** Scatterplot showing the normalized read counts from DESeq2 of well-characterized elongation and paused factors in control, Y14 **(C)** and eIF4AIII **(D)** depleted S2R+ cells. Two of the pre-EJC targets (*MAPK* and *PMCA*) are shown in red as controls. **(E)** RT-qPCR quantification depicting percentage recovery of Mago-bound transcripts in RNA immunoprecipitation with HA tagged Mago, compared to the control-HA tag. Error bars represent the standard deviation from three independent biological replicates. **(F)** RT-qPCR analysis showing the efficiency of Cwc22 knockdown. **(G)** ChIP-qPCR experiments showing recruitment of Mago at the TSS of indicated genes, in control and Cwc22 knockdown conditions, as well after treating the cells with a splicing inhibitor (isoginkgetin) for six hours. Note that Mago recruitment was resistant to Cwc22 KD and only mildly affected with the drug treatment. Error bars indicate the standard deviation from the mean of two biological replicates. **(H)** ChIP-qPCR experiments showing recruitment of pre-EJC components (Mago, Y14, and elF4AIII) at the TSS of indicated genes. Note that Mago recruitment was impaired when transcription was blocked using α-amanitin. Error bars indicate the standard deviation from the mean of two biological replicates.

**Figure S5.**
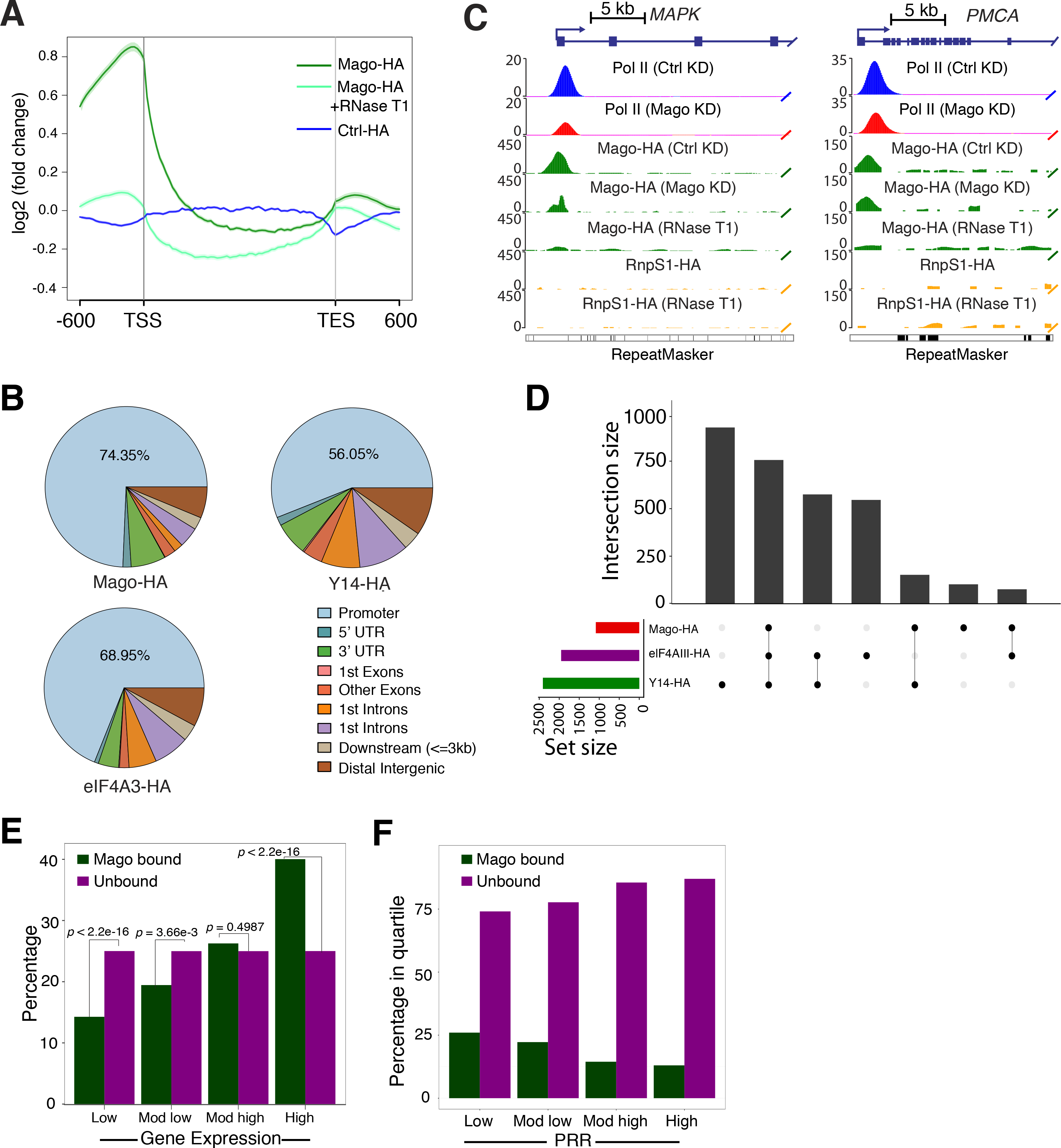
Pre-EJC components are enriched at Promoter Regions of highly expressed and low PRR genes. **(A)** Averaged metagene profile of ChIP-Seq performed with HA-tagged Mago and Ctrl HA-tag in the presence or absence of RNase T1. Log2 fold changes against input control are shown on Y-axis while X-axis depicts scaled genomic coordinates. **(B)** Input normalized and averaged track examples of ChIP-Seq experiments from S2R+ cell extracts transfected with HA-tagged Mago or HA-tagged RnpS1. The cells were either subjected to control or Mago knockdown and chromatins were either untreated or treated with RNase T1, as indicated. Shown here are two pre-EJC target genes *MAPK* and *PMCA*. **(C)** Pie charts showing the distribution of pre-EJC component binding. **(D)** UpSet plot showing the overlap between genes bound by pre-EJC components. The Y-axis shows the size of the intersection while the X-axis shows different conditions in which the overlap was observed, highlighted by line connecting the solid dots. The histogram below to the left depicts number of genes bound by each component. **(E)** Histogram showing percentage of Mago-bound genes amongst different quartiles of genes expression, for genes expressed in control condition. For quartile classification, all of the expressed genes in S2R+ cells were divided into four equal sized quartiles according to the level of their expression, from low to high level. *P*-values for significance of the enrichment for Mago binding amongst different quartiles, derived from Fisher’s exact test, are shown on top of the histogram. **(F)** Histogram showing percentage of genes bound by Mago in different quartiles of PRRs. For quartile classification, all of the expressed genes in S2R+ cells were divided into four equal sized quartiles according to the calculated PRRs, from low to high level. *P*-values for significance of the associations, derived from Fisher’s exact test, are shown on top of the histogram.

**Figure S6.**
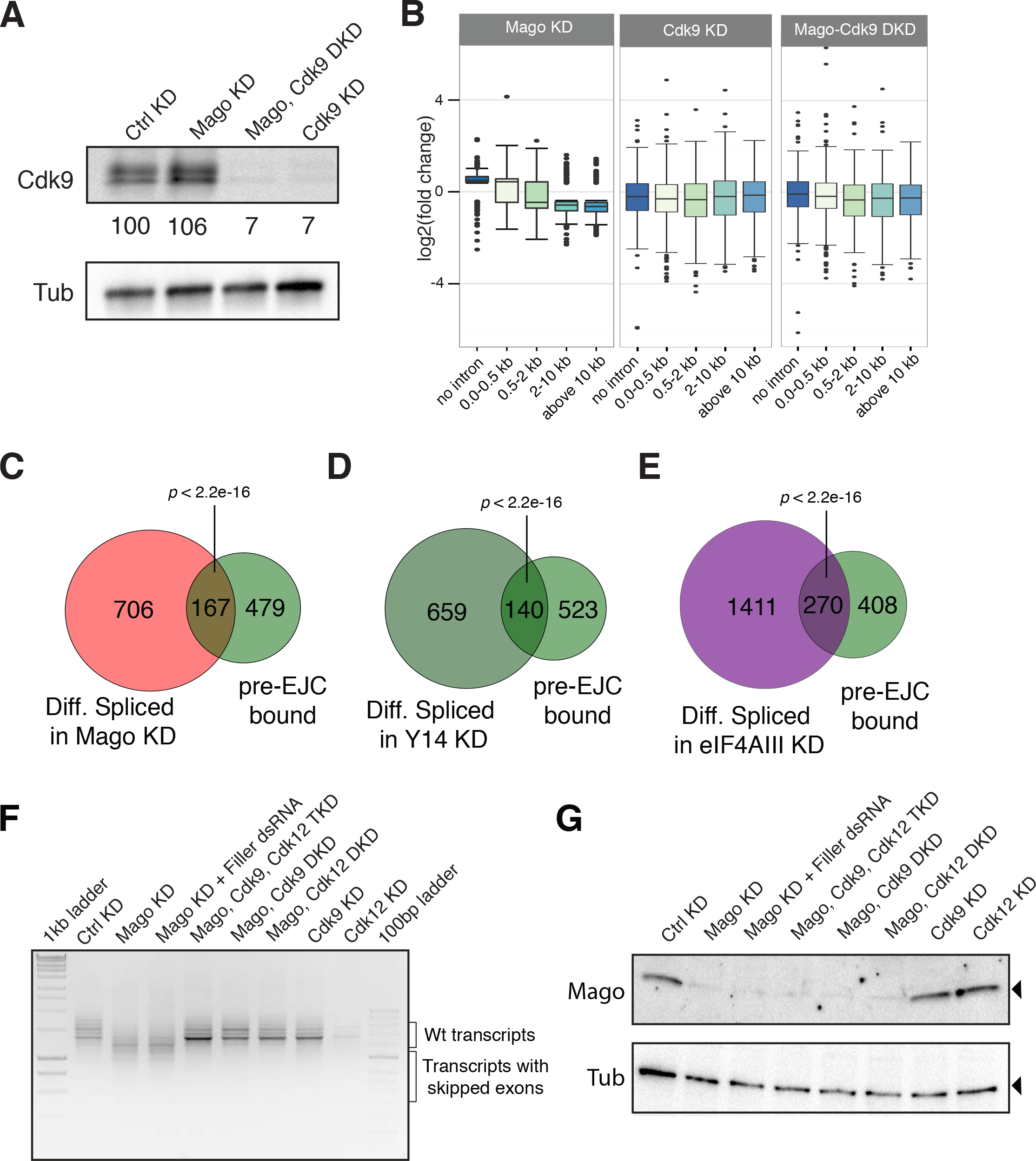
Intron length-dependent changes upon Mago knockdown can by rescued by simultaneous co-depletion of Cdk9 or Cdk12, related to Figure 4 and 6. **(A)** Western blot using antibody directed against Cdk9, showing expression and knockdown efficiency of Cdk9 in S2R+ cells in indicated conditions. Signal in the knockdown conditions was normalized to the control condition using Tubulin as loading control and quantification of the intensity was performed with ImageJ. **(B)** Depletion of Mago in S2R+ cells results in differential gene expression in an intron size dependent manner, when compared to the control condition. Double knockdown of Mago and Cdk9 results in loss of the size dependency effect on gene expression. Shown also is Cdk9 knockdown which affects gene expression independently of intron size, when compared to the control condition. **(C-E)** Venn diagrams showing the overlap between pre-EJC bound genes and genes with differential splicing upon pre-EJC components KD. The overlaps between genes that are differentially spliced upon pre-EJC components KD, and are simultaneously bound by pre-EJC are significant. *P*-values are derived from Fisher’s exact test. **(F)** Agarose gels showing RT-PCR products for *MAPK* using RNA extracted from S2R+ cells with indicated knockdowns as template for cDNA synthesis. The primers used span the 5’ and 3’ UTR of *MAPK*, as described before. Note that similar to Mago and Cdk9 double knockdown, co-depletion of Cdk12 and Mago also rescues *MAPK* splicing defects. The triple knockdown of Mago, Cdk9, and Cdk12 has a slightly better rescue of *MAPK* splicing defects than either of the double knockdowns. **(G)** Western blot using antibody directed against Mago, showing expression and knockdown efficiency of Mago in S2R+ cells in indicated conditions. Shown below is the western blot against Tubulin, used as loading control.

**Figure S7.**
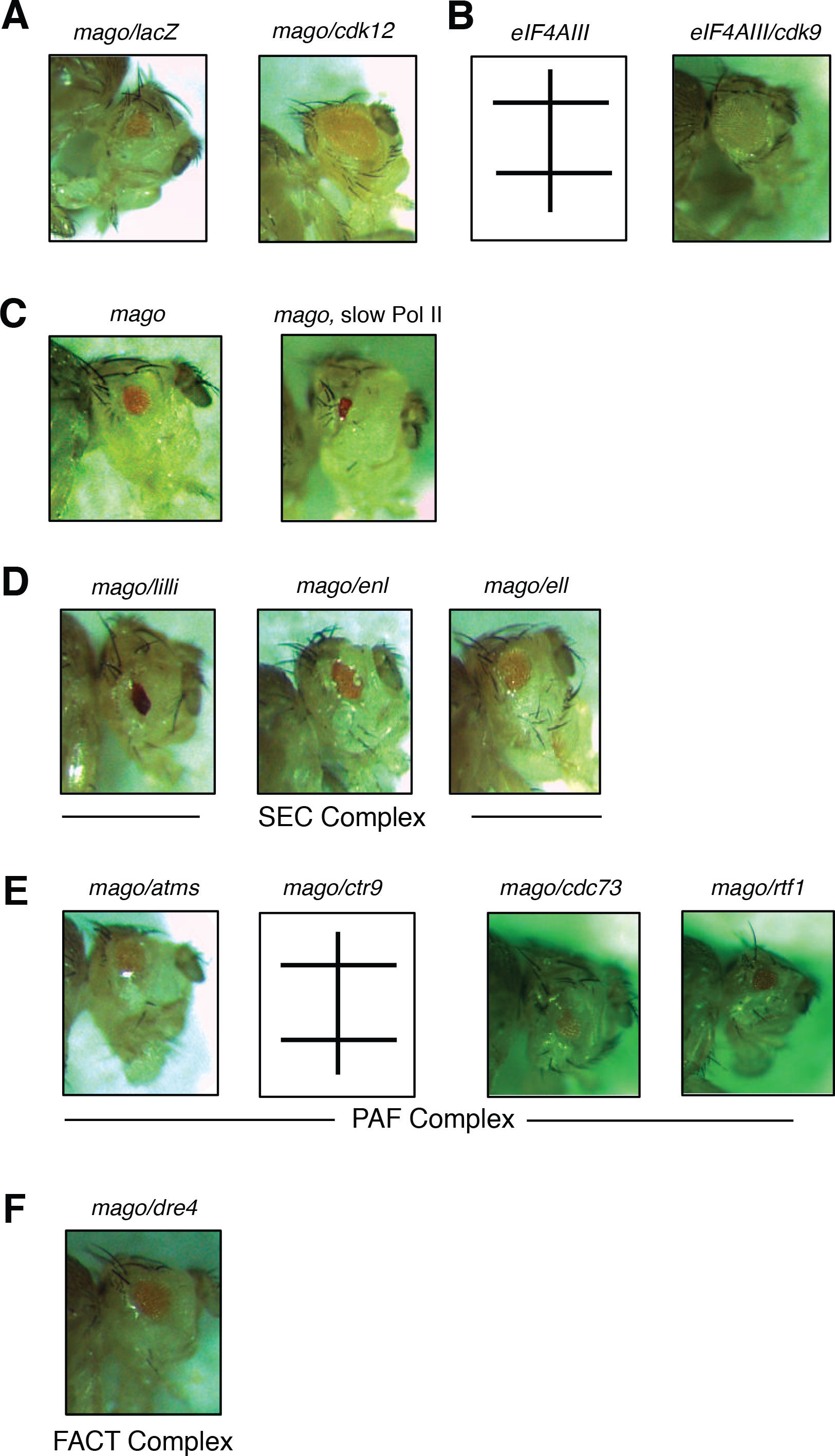
Depletion of Elongation Factors or Slowing Down RNA Pol II Does Not Rescue Mago Photoreceptor Differentiation Defects, related to figure 6. **(A-F)** *Drosophila* adult eyes in different conditions. **(A)** Loss of Mago in the eye results in impairment of eye development due to lack of photoreceptor differentiation. Simultaneous depletion of Cdk12 restores photoreceptor differentiation. **(B)** Loss of eIF4AIII in the eye results in lethality. Simultaneous depletion of Cdk9 restores viability and photoreceptor differentiation. **(C)** Slowing down the kinetics of Pol II using a slow *Pol II* mutant fails to rescue Mago’s effect on photoreceptor differentiation. **(D-F)** Knockdowns of SEC complex components **(D)**, PAF complex components **(E)** or the Dre4 FACT complex subunit **(F)**, do not substantially rescue the eye phenotype resulting from Mago depletion.

## CONTACT FOR REAGENT AND RESOURCE SHARING

As Lead Contact, Jean-Yves Roignant is responsible for all reagents and resource requests. Please contact Jean-Yves Roignant at with requests and inquiries.

## EXPERIMENTAL MODEL AND SUBJECT DETAILS

### *Drosophila* Stocks and Genetics

Fly stocks used were *eyeless*-GAL4, *RpI1215*^*C4*^ (Bloomington *Drosophila* Stock Center). RNAi lines were from the Vienna *Drosophila* Resource Centre (VDRC): Mago (GD28132), Ell (GD34458), DRE4 (GD10916), ENL (GD15671), Lilli (KK106142), Atms (GD20876); and the shRNAi lines from the Transgenic RNAi project (TRIP) and Bloomington *Drosophila* Stock Center: eIF4AIII (32907), Cdk9 (34982), Cdc73 (35238), Rtf1 (36586), Ctr9 (33736).

## METHOD DETAILS

### Cloning

The plasmids used for chromatin immunoprecipitation (ChIP) and co-IP assays in *Drosophila* S2R+ cells were constructed by cloning the corresponding cDNA in the pPAC vector either with N-terminal Flag - 3X HA tag or with HA-SBP tag.

### RNA Isolation and RT-PCR

RNA was extracted from cells using Trizol reagent, following the manufacturer’s protocol. For reverse transcription, cDNA was synthesized using MMLV reverse transcriptase (Promega, Cat No-M1701). For semi-quantitative RT-PCR 2 μg of RNA was reverse transcribed. 5 μ1 of the cDNA was amplified using the respective primers in 50 μ1 PCR reaction, using One Taq polymerase (NEB, Cat No-M0480). After 40 cycles of amplification half of the PCR product was loaded on 1 % agarose gel to qualitatively analyze the splicing products. For real time PCR, Rpl15 was used as an internal control. Relative abundance of transcripts was calculated by the 2^Δ^ Ct method. PCR primers used for semi-quantitative and real time PCR are listed in Table 1.

**Table 1.**
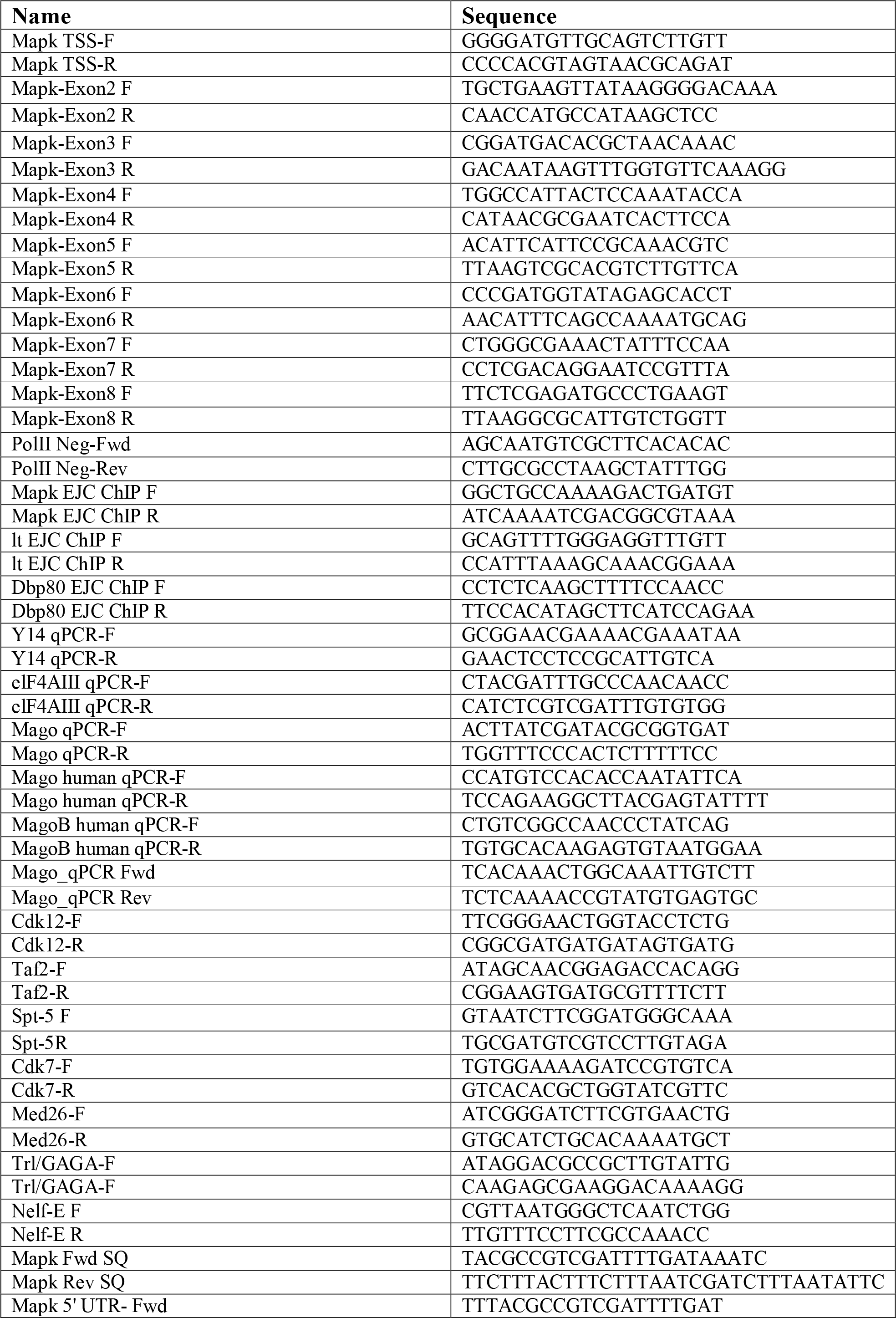
Sequences of the primers used for qRT-PCR and semi-quantitative RT-PCR

STAR □ Methods
| REAGENT or RESOURCE | SOURCE | IDENTIFIER |
| --- | --- | --- |
| **Antibodies** |  |  |
| Total Pol II | Diagenode | Cat No-15200004 |
| Ser2P | Abcam | Ab5095 |
| Flag-M2 | Sigma | Cat No-F3165 |
| Ser5P | Chromotek | 3E8 |
| Ser2P | Chromotek | 3E10 |
| hCdk9 | Santa Cruz | Cat No-8338 |
| Polyclonal against dMago | Metabion | Custom made |
| Magoh | Santa Cruz | Cat No- 56724 |
| **Bacterial and Virus Strains** |  |  |
| DH5α | | |
| Biological Samples |  |  |
| **Fly stocks** | Bloomington *Drosophila* Stock Center, the Transgenic RNAi Project in Harvard and the Vienna *Drosophila* Resource Center |  |
| **Chemicals, Peptides, and Recombinant Proteins** |  |  |
| MMLV reverse transcriptase | Promega | Cat No-M1701 |
| One Taq polymerase | NEB | Cat No-M0480 |
| Schneider Cell's Medium | GIBCO | Cat No-21720 |
| Effectene Transfection Reagent | Qiagen | Cat No-301425 |
| Hi-Scribe T7 transcription kit | NEB | Cat No-E2040 |
| RNAiMax | Invitrogen | Cat No-13778030 |
| NebNext Ultra II DNA library kit | NEB | Cat No- E7645 |
| True Seq stranded mRNA preparation kit | illumina | Cat No- 20020594 |
| 5,6-dichlorobenzimidazole 1-β-d-ribofuranoside | Sigma | D1916 |
| 4- thiouridine | Sigma | T4509 |
| EZ Link Biotin HDPD | Thermo Fisher | Cat No-21314 |
| Critical Commercial Assays |  |  |
| SuperSignal™ West Pico Chemiluminescent Substrate | Thermo Scientific | Cat No-34080 |
| **Deposited Data** |  |  |
| ChIP-Seq, RNA-Seq, 4sU-seq, MNase-seq and DamID-seq | GEO | Superseries- 92389 |
| **Experimental Models** |  |  |
| HeLa cells |  |  |
| S2R+ cells | DGRC |  |
| **Experimental Models: Organisms/Strains** |  |  |
| *Drosophila Melanogaster* |  |  |
| **Oligonucleotides** |  |  |
| All the primers used in this study | Table -1 |  |
| **Recombinant DNA** |  |  |
| pA5.1C-Mago-Flag-HA |  |  |
| pA5.1C-Cdk9-SBP-HA |  |  |
| pA5.1C-Rnps1-Flag-HA |  |  |
| pA5.1C-Y14-Flag-HA |  |  |
| pA5.1C-eIF4AIII-Flag-HA |  |  |
| pA5.1C-BoxB-FLUC |  |  |
| pA5.C-Renilla |  |  |
| pA5.1C-BoxB |  |  |
| pA5.1C-lambdaN-Mago |  |  |
| pA5.1C-lambdaN-RnpS1 |  |  |
| pA5.1C-lambdaN-eGFP |  |  |
| pA5.1C-lambdaN |  |  |
| **Software and Algorithms** |  |  |
| STAR | v2.5.1b |  |
| htseq Count | v0.6.1p1 |  |
| DeSeq2 | v1.10.1 |  |
| rMATS | v3.2.1b |  |
| MACS2 | v2.1.1 |  |
| bowtie2 | v2.2.8 |  |
| deepTools | v2.2.3 |  |
| NGS.plot | v2.61 |  |
| R | v3.4.2 |  |

### Cell Culture, RNAi and Transfection

*Drosophila* S2R+ cells were cultured in Schneider Cell’s Medium (GIBCO, Cat No-21720) supplemented with 10% FBS and 2% Penicillin/Streptomycin. The plasmids expressing various transgenes were transfected with Effectene transfection reagent (Qiagen, Cat No- 301425), following manufacturer’s protocol. For knock down experiments, dsRNA was synthesized overnight at 37 °C using Hi-Scribe T7 transcription kit (NEB, Cat No-E2040). dsRNA was transfected in S2R+ cells by serum starvation for 6 hours. The treatment was repeated three times and cells were harvested 7 days after the first treatment for Mago. For knockdown of other pre-EJC components and Btz, the treatment was repeated two times and cells were harvested 5 days after the first treatment. The primers used for generating dsRNA are listed on Table S2. S2R+ Cells were treated with 50 μg/ml of α-amanitin for 7 hours to block transcription.

HeLa cells were cultured in standard RPMI medium supplemented with 10% FBS and 2% Penicillin/Streptomycin. For siRNA knockdown, cells were transfected with 10 nM of siRNA using RNAiMax (Invitrogen) according to manufacturer’s protocol. Cells were harvested 48 h after transfection. A mixture of three siRNA was used to deplete Magoh, two against MagohA isoform (siRNA sequence; 1-CGGGAAGTTAAGATATGCCAA; 2-CAGGCTGTTTGTATATTTAAT) and one targeting MagohB isoform (siRNA sequence; GATATGCCAACAACAGCAA).

### Antibodies

The following antibodies were used in this study. For total Pol II ChIP, RBP1 (Diagenode Cat No-15200004) was used. Ser2P ChIP was performed using ab5095 (Abcam); 3E10 (Chromotek) was used for western blotting. Anti-Ser5P Pol II (Chromotek, Cat No-3E8) and ARNA3 (Pol II) antibodies (Progen, Cat No-65123) were used for western blot assays. For immunoprecipitation experiments, anti-Flag M2 (Sigma, Cat No-F3165) and M-280 streptavidin beads (ThermoFisher, Cat No-11205) were used. A polyclonal rabbit anti-Mago antibody was generated from Metabion (Germany). Anti-hCdk9 (Santa Cruz, Cat No-8338) was used for immunoprecipitation from HeLa cells extracts. The Cdk9 western blot from S2R+ cell extract was performed by anti-dCdk9 antibody, a kind gift from Akira Nakamura.

### Immnunostaining

The primary antibodies used were rat anti-Elav (1:5; Developmental Studies Hybridoma Bank) and guinea pig anti-Senseless (1:1000) (Frankfort et al., 2001). Eye imaginal discs were dissected in 0.1 M sodium phosphate buffer (pH 7.2) and then fixed in PEM (0.1 M PIPES at pH 7.2 mM MgSO4, 1 mM EGTA) containing 4% formaldehyde. Washes were done in 0.1 M phosphate buffer with 0.2% Triton X-100. Appropriate fluorescent-conjugated secondary antibodies were used (1:1000; Jackson Immunoresearch Laboratories). Images were collected on Zeiss TCS SP5 confocal microscope.

### Co-IP Assay and Western Blot Analysis

For co-IP assay in S2R+ cells, cells were plated in 10 cm cell culture dish, and respective transgenes were transfected using Effectene transfection reagent, according to manufacturer’s protocol. 48h post transfection cells were collected, washed once with PBS and re-suspended in swelling buffer (10mM Tris pH 7.5, 2 mM MgCl2, 5 mM MgCl2, 3mM CaCl2, and protease inhibitors). After incubating 10 min on ice, the suspension was spun at 600g for 10 min at 4°C. After discarding the supernatant the pellet was resuspended in lysis buffer (10mM Tris pH 7.5, 2 mM MgCl2, 5 mM MgCl2, 3mM CaCl2, 0.5% NP-40, 10% glycerol and protease inhibitors) and centrifuged for 5 min at 600g. Nuclei were resuspended in lysis buffer (40mM HEPES pH7.4, 140mM NaCl, 10 mM MgCl2, 0.5% Triton X-100 and protease inhibitors) and sonicated in the bioruptor plus (Diagenode) for 6 cycles with 30 Sec “ON/OFF” at low settings. Protein concentrations were determined using Bradford reagent (BioRad, Cat No-5000006). For IP 2 mg of proteins were incubated with respective antibody in lysis buffer and rotated head-over-tail O/N at 4 °C. The beads were washed 3x for 10 min with lysis buffer and IP proteins were eluted by incubation in 1x SDS buffer at 85 °C for 10 min. Immunoprecipitated and input proteins were analyzed by western blot, after separating them on 4-15% gradient SDS-PAGE gel (BioRad, Cat No-4561083) and transferred to PVDF membrane (Millipore, Cat No-IPVH00010). After blocking with 5 % milk in TBST (0,05 % Tween in 10mM Tris pH 7.4 and 140mM NaCl) for O/N at 4 °C, the membrane was incubated with respective primary antibody in blocking solution O/N at 4 °C. Membrane was washed 4 × in TBST for 15 min and incubated 1 h at RT with secondary antibody in blocking solution. Blots were developed using SuperSignal™ West Pico Chemiluminescent Substrate (Thermo Scientific, Cat No-34080) and visualized using BioRad Gel documentation system.

For co-IP assay in HeLa, cells were plated in 10 cm cell culture dish. Afterwards, cells were collected washed once with PBS and re-suspended in swelling buffer (10mM Tris pH 7.5, 2 mM MgCl2, 5 mM MgCl2, 3mM CaCl2, and protease inhibitors), identical to the approach as in S2R+ cells. After incubating 10 min on ice, the suspension was spun at 600g for 10 min at 4°C. After discarding the supernatant the pellet was resuspended in lysis buffer (10mM Tris pH 7.5, 2 mM MgCl2, 5 mM MgCl2, 3mM CaCl2, 0.5% NP-40, 10% glycerol and protease inhibitors) and centrifuged for 5 min at 600g. Nuclei were resuspended in lysis buffer (40mM HEPES pH7.4, 140mM NaCl, 10 mM MgCl2, 0.5% Triton X-100 and protease inhibitors) and sonicated in the bioruptor plus (Diagenode) for 6 cycles with 30 Sec “ON/OFF” at low settings. Protein concentrations were determined using Bradford reagent (BioRad, Cat No-5000006). For IP 2 mg of proteins were incubated with anti-Magoh antibody in lysis buffer and rotated head-over-tail O/N at 4 °C. The beads were washed 3x for 10 min with lysis buffer and IP proteins were eluted by incubation in 1x SDS buffer at 85 °C for 10 min, immunoprecipitated and input proteins were analyzed by western blot, as described above.

### RNA extraction and RNA-seq

RNA was extracted from cells using Trizol reagent, following the manufacturer’s protocol. RNA was further cleaned for organic contaminants by RNeasy MinElute Spin columns (Qiagen, Cat No-74204). The purified RNA was subjected to oligodT (NEB, Cat No-S1419S) selection to isolate mRNA. The resulting mRNA was fragmented and converted into libraries using illumina TruSeq Stranded mRNA Library Prep kit (illumina, Cat No-20020594) following manufacturer’s protocol.

### ChIP-qPCR and ChIP-Seq

S2R+ cells and HeLa cells were fixed with 1% formaldehyde for 10 min at room temperature, and harvested in SDS buffer resuspended in RIPA buffer (140 mM NaCl, 10 mM Tris-HCl [pH 8.0], 1 mM EDTA, 1% Triton X-100, 0.1% SDS, 0.1% DOC), and lysed by sonication. The lysate was cleared by centrifugation, and incubated with respective antibodies overnight at 4°C. Antibody complexes bound to protein G beads were washed once with 140 mM RIPA, four times with 500 mM RIPA, once with LiCl buffer and twice with TE buffer for 10 min each at 4°C. DNA was recovered after reverse crosslinking and phenol chloroform extraction. After precipitating and pelleting, DNA was dissolved in 30 μ1 of TE. Control immunoprecipitations were done in parallel with either tag alone or knock down controls, and processed identically. 5 μ1 of immunoprecipitated DNA was used for checking enrichment with various primer pairs (listed in supplementary table 1) on Applied Biosystem ViiA™ 7 real time machine using SYBR green reagent (Life technologies, Cat No-4367659). After validating enrichment, the recovered DNA was converted into libraries using NebNext Ultra DNA library preparation kit, following manufacturer’s protocol. DNA libraries were multiplexed, pooled and sequenced on Illumina HiSeq 2000 platform.

### DRB-4sU-Seq

S2R+ cells were grown in Schneider’s Cell Medium with 10% bovine serum supplemented with antibiotics and maintained at 25°C. 5,6-dichlorobenzimidazole 1-β-d-ribofuranoside (DRB) from Sigma (D1916) was used at a final concentration of 300 μM, dissolved in water, for 5 h. 4-thiouridine (4sU) was purchased from Sigma (Cat No-T4509) and used at a final concentration of 100 μM. Control and Mago KD was performed as described before. All the samples were labeled for 6 min with 4-thiouridine, and transcription was allowed to proceed after DRB removal for 0, 2, 8 and 16 min along with one non-DRB treated control.

A total of 100 to 130 μg RNA was used for the biotinylation reaction. 4sU-labeled RNA was biotinylated with EZ-Link Biotin-HPDP (Thermo Scientific, Cat No-21341), dissolved in dimethylformamide (DMF, Sigma Cat No-D4551) at a concentration of 1 mg/mL. Biotinylation was done in labeling buffer (10 mM Tris pH 7.4, 1 mM EDTA) and 0.2 mg/mL Biotin-HPDP for 2 h with rotation at room temperature. Two rounds of chloroform extractions removed unbound Biotin-HPDP. RNA was precipitated at 20,000 g for 20 min at 4°C with a 1:10 volume of 5M NaCl and an equal volume of isopropanol. The pellet was washed with 75% ethanol and precipitated again at 20,000 g for 10 min at 4°C. The pellet was left to dry, followed by resuspension in 100 μL RNase-free water. Biotinylated RNA was captured using Dynabeads MyOne Streptavidin T1 beads (Invitrogen, Cat No-65601). Biotinylated RNA was incubated with 50 μL Dynabeads with rotation for 15 min at 25°C. Beads were magnetically fixed and washed with 3× Dynabeads washing buffer. RNA-4sU was eluted with 100 μL of freshly prepared 100 mM dithiothreitol (DTT), and cleaned on RNeasy MinElute Spin columns (Qiagen, Cat No-74204). For the untreated 4sU-Seq version used for calculating polymerase release ratio (PRR), an identical approach was used with following modifications. During the period when biotinylated RNA was incubated with 50 μL Dynabeads with rotation for 15 min at 25°C, RNAse T1 was added in order to fragment RNA to 100bp. Beads were magnetically fixed and washed with 3× Dynabeads washing buffer, as described before. RNA-4sU was eluted with 100 μL of freshly prepared 100 mM DTT, and cleaned on RNeasy MinElute Spin columns (Qiagen, Cat No-74204). Enriched nascent RNAs were converted to cDNA libraries with *Drosophila* Ovation Kit (Nugen- Cat No-7102-32) with integrated ribosomal depletion workflow. Amplified cDNA libraries were pooled, multiplexed, and sequenced on two lanes of Illumina HiSeq 2000.

### MNase-Seq

S2R+ cells were fixed with 1% formaldehyde for 10 min at RT. Cells were harvested and 20 million nuclei were spun at 3,500 g at 4°C for 10 min. Nuclear Pellet was resuspended in 300 μl of MNase digestion buffer (0.5 mM spermidine, 0.075% NP40, 50 mM NaCl, 10mM Tris-HCl, pH 7.5, 5mM MgCl2, 1mM CaCl2, 1mM β◻mercaptoethanol and complete protease inhibitors). Reaction was spun at 3,200g, 4°C for 10 min and resuspended in 50 μl of MNase digestion buffer and digested with 30U of MNase at 37°C for 10 min at 300 rpm in mixing block. The MNase digestion reaction was quenched with EDTA at 10 mM final concentration. After 10 min on ice, the nuclei were washed once with 1 ml of RIPA buffer (140 mM NaCl and complete protease inhibitors). Pellet was resuspended in 300 μl of RIPA buffer (140mM) and sonicated (3 cycles, medium intensity, 30 s on/off intervals) and centrifuged at 18,000g, 4°C for 10 min. DNA was recovered after reverse crosslinking and phenol chloroform extraction. After precipitating and pelleting, DNA was dissolved in 30 μl of TE and resolved on agarose gel. The ~ 147bp fragments corresponding to the mono nucleosomal fragments were gel extracted and subjected to 50 bp paired end sequencing on Illumina HiSeq 2500 platform.

### DamID-Seq

pUAST-LT3-ORF1 vector (kind gift from A. Brand) were used to clone Cdk9 as a C-terminal Dam-fusion protein. The Dam-Cdk9 itself was cloned downstream of mcherry (as a primary ORF) separated by stop codon. This ensured low level expression of the Dam-Cdk9 fusion protein. S2R+ cells were plated in 10 cm dish and subjected to control and Mago knockdowns using dsRNA, as described earlier. On the sixth day of knockdown pUAST-LT3-Dam-Cdk9 was co-transfected with pActin-Gal4 vector to induce Dam-Cdk9 expression, using effectene transfection reagent according to manufacturer’s protocol. The Dam alone control was similarly transfected in control and Mago depleted S2R+ cells. DNA was isolated from cells after 16 h of transfection and subsequent treatments were performed as described (Marshall et al., 2016). Purified and processed genomic DNA of two biological duplicates was subjected to library preparation using the NebNext DNA Ultra II library kit (New England Biolabs) and sequenced on a NextSeq500.

### Computational Analysis

RNA-Seq: Libraries of strand-specific RNA-seq were constructed using Illumina TruSeq Kit, following the manufacturer’s protocol. The libraries were sequenced with a read length of 71 bp in paired end mode. Mapping was performed using STAR (Dobin et al., 2013)(v. 2.5.1b). Counts per gene were derived using htseq count (v.0.6.1p1). Differential expression analysis was done using DESeq2 (Love et al., 2014)(v.1.10.1), differential expressed genes were filtered for an FDR of 1% and a fold change of 1.3. Splicing analysis was done using DEXSeq (Anders et al., 2012)(v. 1.16.10) with 10% FDR filtering, and rMATS (Shen et al., 2014)(v. 3.2.1b) with 10% FDR. Genes were defined as expressed if they had coverage above 1 rpkm. ChIP-Seq: The libraries were sequenced on a HiSeq2500 in either paired end or single end mode. De-multiplexing and fastq file conversion was performed using blc2fastq (v.1.8.4). Libraries (Ser2P ChIP, RnpS1, and Control Pol II) were de-multiplexed using 6 bp front tags. After sorting, the tags and the A-overhang base were trimmed (7 bp in total). The other libraries were demultiplexed with inline adaptors. Reads from ChIP-Seq libraries were mapped using bowtie2 (Langmead and Salzberg, 2012)(v. 2.2.8), and filtered for uniquely mapped reads. The genome build and annotation used for all *Drosophila* samples was BDGP6 (ensemble release 84). The genome build and annotation used for the HeLa samples was hg38 (ensemble release 84). Peak calling was performed using macs2 (Zhang et al., 2008)(v 2.1.1-20160309). Further processing was done using R and Bioconductor packages. Input normalized bigwig tracks were produced using Deeptools (Ramirez et al., 2016)(v. 2.2.3). Heatmaps and input normalized tracks were produced using Deeptools (v. 2.2.3). Metagene profiles were produced using NGS.plot (v. 2.61) and Deeptools (v. 2.2.3) after input and “spike-in” normalization.

To assign the target genes bound by pre-EJC components, peaks were called using MACS2 with 2.0 fold enrichment as cut-off. The resulting peaks were annotated with the ChIPseeker package on Bioconductor, using nearest gene to the peak summit as assignment criteria. The intersection of genes bound by all pre-EJC components, i.e Mago-HA, Y14-HA, and eIF4A3-HA, was defined as pre-EJC bound.

4sU-Seq: The libraries were sequenced with a read length of 50 bp in single end mode. Mapping was performed using STAR (v. 2.5.1b). Reads mapping to the exons were removed using annotation (ENSEMBL v73) for *Drosophila*. Multimapped reads were filtered out, and uniquely mapped reads to the transcript was considered.

Calculation of Polymerase Release Ratio (PRR): Polymerase release ratios (PRRs) were calculated as follows: for each gene, the TSS region was defined as 250 bp upstream to 250 bp downstream of the TSS. The gene body was defined as 500 bp downstream of the TSS to 500 bp upstream of the TES. The PRR ratio was calculated as the log2 ratio between the enrichment in the downstream region towards the enrichment at the TSS. For each gene, the TSS with the highest average signal around the TSS in the Control condition was selected. Enrichment calculations were based on the enrichment over the input (ChIP-Seq) or t0 (4sU-Seq).

Calculation of Elongation Rate: For elongation rate calculation, all the genes longer than 10 kb were divided into 100 bp bins (to a total of 20 kb) and the transcriptional wave front was identified in the bin with lowest local minima signal. The distance covered by the wave front between 2 min after DRB removal and 8 min is then divided by the corresponding time interval of 6 min to calculate elongation rates.

MNase-Seq: The libraries were sequenced with a read length of 50 bp in paired end mode. Demultiplexing and fastq file conversion was performed using blc2fastq (v.1.8.4). Libraries were de-multiplexed using 6 bp front tags. After sorting, the tags and the A-overhang base were trimmed (7 bp in total). Reads from MNase-Seq libraries were mapped using bowtie2 (v. 2.2.8), and filtered for uniquely mapped reads. The genome build and annotation used for all *Drosophila* samples was BDGP6 (ensemble release 84). Further processing was done using R and Bioconductor packages. Heatmaps and input normalized tracks were produced using Deeptools (v. 2.2.3). Metagene profiles were produced using NGS.plot (v. 2.61).

Targeted DamID-Seq: The libraries were sequenced with a read length of 50 bp in paired end mode. The first read was mapped to Drosophila melanogaster genome (BDGP6) using bowtie (v.2.2.9), binned to GATC fragments, and normalized against the Dam-only control (Marshall and Brand, 2015) using the available damidseq_pipeline on GitHub. The resulting bedgraph files were averaged and smoothened using BEDOPS (v. 2.4.30). The smoothened bedgraph files were converted to bigwig file using SeqPlot, and processed through Deeptools (v. 2.2.3) to generate heatmaps. To quantify the changes at the promoter, the signals in the bedgraph were mapped to the promoters using bedmap tool available in BEDOPS software. The further quantification and plots were generated using R (v 3.4.2), and ggplot2 package available on Bioconductor.

## QUANTIFICATION AND STASTISTICAL ANALYSIS

Statistical parameters and significance are reported in the Figures and the Figure legends. For comparisons of the distribution of different classes we used ANOVA. t-Test, two-sample Kolmogorov-Smirnov test and Fisher’s test were used for testing the statistical significance.

